# DNA Origami – Lipid Membrane Interactions Defined at Single-Molecular Resolution

**DOI:** 10.1101/2023.11.14.567022

**Authors:** Elena Georgiou, Javier Cabello-Garcia, Yongzheng Xing, Stefan Howorka

## Abstract

Rigid DNA nanostructures that bind to floppy bilayer membranes are of fundamental interest as they replicate biological cytoskeletons for synthetic biology, biosensing, and biological research. Here, we establish principles underpinning the controlled interaction of DNA structures and lipid bilayers. As membrane anchors mediate interaction, more than 20 versions of a core DNA nanostructure are built each carrying up to five individual cholesterol anchors of different steric accessibility within the 3D geometry. The structures’ binding to membrane vesicles of tunable curvature is determined with ensemble methods and by single-molecule localization microscopy. This screen yields quantitative and unexpected insight on which steric anchor points cause efficient binding. Strikingly, defined nanostructures with a single molecular anchor discriminate effectively between vesicles of different nanoscale curvatures which may be exploited to discern diagnostically relevant membrane vesicles based on size. Furthermore, we reveal anchor-mediated bilayer interaction to be co-controlled by non-lipidated DNA regions and localized membrane curvatures stemming from heterogenous lipid composition, which modifies existing biophysical models. Our study extends DNA nanotechnology to control interactions with bilayer membranes and thereby facilitate the design of nanodevices for vesicle-based diagnostics, biosensing, and protocells.

---

Defined DNA nanostructures that bind to floppy semifluid bilayer membranes are scientifically intriguing and innovatively combine the best of DNA nanotechnology and membranes for applications in synthetic biology, biosensing, and research into cell biology, biophysics, and biomimetics. While DNA nanotechnology excels at precisely tuning the shape and dimensions of 2D and 3D nanostructures spanning from the nanoscale^1–13^ to the macroscale,^13–15^ semifluid lipid bilayers stand out by compartmentalizing hydrophobic environments,^16^ setting up concentration gradients for energy conversion,^17^ and providing lateral diffusive platforms for enhanced molecular assembly.^18^ Combining DNA nanotechnology with lipid membranes can unlock considerable synergy as illustrated by a range of DNA nanostructures that can mimic cellular cytoskeletons and shape membranes into biologically unprecedented forms,^19–23^ define lipid domains,^24^ selectively label leaflets,^25, 26^ measure membrane curvature,^27^ fuse membrane bilayers,^28^ spatially activate membrane proteins,^29, 30^ tune endosomal uptake,^31^ facilitate macrostructure assembly,^32, 33^ and even help produce synthetic protocells.^34–37^ In complementary approaches, DNA nanostructures can insert into lipid membranes to emulate the function of membrane proteins, including receptors,^38^ nanopores,^39, 40^ gated channels,^41^ membrane force sensors,^42^ and lipid flippases.^43^ Designing bilayer-interacting DNA nanostructures hinges on attached hydrophobic anchors that insert into the lipid bilayer.^44, 45^ Cholesterol is the most prominent anchor, yet tocopherol,^46^ porphyrins,^47^ alkyl chains,^45^ and polypropylene oxide have also been successfully used.^48^

Understanding DNA-membrane interaction is of fundamental scientific relevance but also helps improve the engineering of DNA nanostructures. Several studies have investigated how cholesterol-mediated anchoring depends on temperature, lipid composition, the number of cholesterol anchors, and buffer conditions.^49–55^ However, very few studies^49, 54^ have explored the impact of steric and geometric effects even though they are a principal contributor to ligand-binding specificity in both chemistry and biology^56–58^ and are likely key for controlling DNA nanostructure binding to bilayer. Unresolved questions are how the nanostructure-membrane interactions depend on the steric accessibility, position, and number of anchors in a DNA nanostructure of 3D geometry. A related question is on the role of the membrane geometry in terms of global vesicle curvature but also smaller localized curvatures which can result from non-homogenous lipid composition.^59^ Ideally, these questions should be addressed with ensemble techniques for efficient throughput but also at the single-molecule level to obtain further insight into the binding mechanism. A comprehensive understanding of steric influence would be of scientific value and help identify design rules for efficient membrane binding and nanostructure engineering for size-specific vesicle discrimination to identify diagnostically relevant exosomes.^60, 61^

Here, we comprehensively examine how steric factors influence DNA nanostructure-membrane interactions. We devise a T-shaped DNA nanoprobe (DNP) structure with prominent geometry featuring a baseplate and a tip (Figure 1A). DNP is a highly addressable 3D breadboard to place cholesterols of variable steric accessibility. We use 20 different DNA variants (Figure 1B) which are clustered into groups 1-4, each comprising up to five cholesterol tags (Figure 1B, DNP-1 to DNP-4, top row; Figure S1). In groups DNP-1 and DNP-2, cholesterol moieties are positioned at one corner of the baseplate (Figure 1B, insets) and likely of intermediate to high steric accessibility for vesicles (Figure S1, S2). By contrast, lipidated DNP variants of group 3 have cholesterols very close to or on the DNP tip (Figure 1B, Figure S1), likely of lower accessibility (Figure S2), while group DNP-4 structures feature cholesterols at the center of the bottom baseplate distant to corner and the tip (Figure 1B, Figure S2). Predicted accessibility also guides the positioning of cholesterols within the groups, such as in DNP-1 where the first cholesterol is close to the corner while the second and third cholesterols are closer to the tip in the middle of the baseplate (Figure 1B, middle row). The lipidated nanostructure variants are termed DNP-1.1 to DNP-1.4 whereby the second digit refers to the number of cholesterols (Figure 1B, middle row, Figure S1). DNP variants in group 2-4 with one to up to five cholesterols are named accordingly.

**Figure 1.**
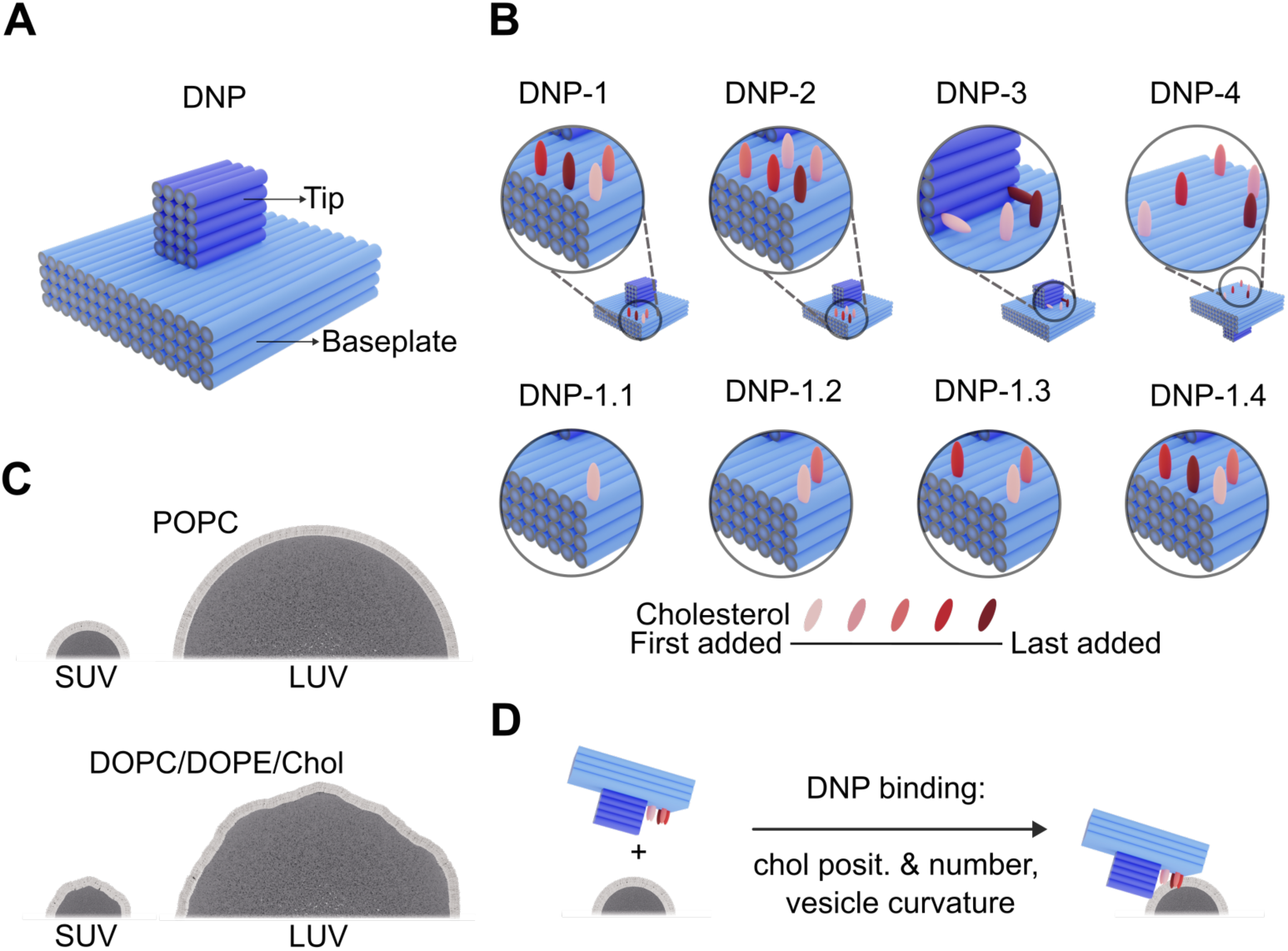
DNA nanoprobes (DNPs) and their interaction with bilayer vesicles as a function of membrane anchor number, anchor position, and vesicle curvature and composition. (A) Design and dimensions of the T-shaped DNP with baseplate (light blue) and tip (dark blue) made of bundled DNA duplexes (blue cylinders). (B) Lipidated DNP versions of group DNP-1 and DNP-2 carry up to five cholesterols (pink to red) at the edge-corner of the baseplate, while those of DNP-3 at the tip and DNP-4 at the baseplate underside, respectively (top row). Variants of the first group DNP-1.1 to DNP-1.4 carry one to four cholesterols (middle row). Additional cholesterols added within a DNP group are color-coded in red of increasing intensity (bottom row). (C) Schematic illustration of small unilamellar vesicles (SUVs) and large unilamellar vesicles (LUVs) of diameters of 30 nm and 200 nm, and membrane compositions of either POPC with homogeneously curved membrane, or DOPC/DOPE/Chol (2:1:2) with heterogeneous phases of varying localized curvature. (D) Scheme on lipid anchor-mediated binding of DNPs to vesicles. Successful binding is detected with agarose gel electrophoresis and high-resolution single-molecule fluorescence microscopy.

The binding of each of these DNA structures is tested with four lipid vesicle types of different global diameters and more fine-grained local curvature (Figure 1C). We use large unilamellar vesicles (LUV) and small unilamellar vesicles (SUV) of around 200 and 30 nm diameter each made with either 1-palmitoyl-2-oleoyl-*sn*-glycero-3-phosphocholine (POPC)(Table S1) to achieve constant membrane curvature or lipid mixture 1,2-dioleoyl-sn-glycero-3-phosphocholine (DOPC), 1,2-dioleoyl-*sn*-glycero-3-phosphoethanolamine (DOPE), and cholesterol (Chol, ratio of 2:1:2)(Table S1) with heterogeneous phases of varying localized curvatures.^59, 62–64^ To screen for binding, ensemble measurements and single molecule localization microscopy are used and report on which steric factors of DNP and membrane vesicles influence binding yield (Figure 1D).

Our results are multifaceted and (i) quantify how binding correlates with higher cholesterol number on DNPs while revealing that binding strongly depends on the position of the cholesterols, often against previously assumed knowledge. As other striking result, (ii) a selected DNA nanostructure with solely one cholesterol shows strong size selection for vesicles larger than 100 nm while interacting poorly with small vesicles. Based on this and related findings we expand the current model that explains DNA-membrane interactions as mainly governed by lipid anchors to include interactions between non-modified DNA surfaces and other bilayer segments. The surprising finding may help size-separate diagnostically relevant exosome membrane vesicles and thereby fill a gap in the diagnostic toolset. Exosomes are extracellular 30-150 nm-sized vesicles present in most biological fluids which mediate intercellular communication and signaling^65, 66^ and are hence important biomarkers for cancer as well as cardiovascular and neurodegenerative diseases.^67, 68^ As last finding, (iii) homogenous vs heterogenous fine-grained vesicle curvatures strongly influence nanostructure binding, partly more than overall vesicle curvature. By improving understanding of DNA nanostructure binding to membranes, our study may facilitate rational design for synthetic biology, biophysical research, and the purification of diagnostically relevant vesicles.

## RESULTS AND DISCUSSION

### Design, assembly, and characterization of the T-shaped DNA nanostructure

The DNP structure was rationally designed with the software CaDNAno.^69^ In the defined T-shaped DNP, the baseplate is composed of four stacked layers of 16 parallel duplexes arranged in a square lattice whereby each duplex is 79 base pair (bp) long (Figure 1A, Figure 2A, Figure S3). The corresponding dimensions for length, width, and height of the baseplate are predicted to be 26.9 nm x 41.6 nm x 10.4 nm, assuming a duplex length of 0.34 nm per bp and a duplex-duplex distance in the square lattice of 2.6 nm.^70^ At two sides of the baseplate, small few-nucleotide short loops from the scaffold were allowed at each duplex end to prevent aggregation by blunt-end stacking interactions.^1, 71^ The cuboid tip of the nanostructure was designed to be 4 x 5 duplexes each with 31 bp (Figure 2A, Figure S3), corresponding to length, width, and height dimensions of 10.5 nm, 10.4 nm, and 13.0 nm, respectively.

**Figure 2.**
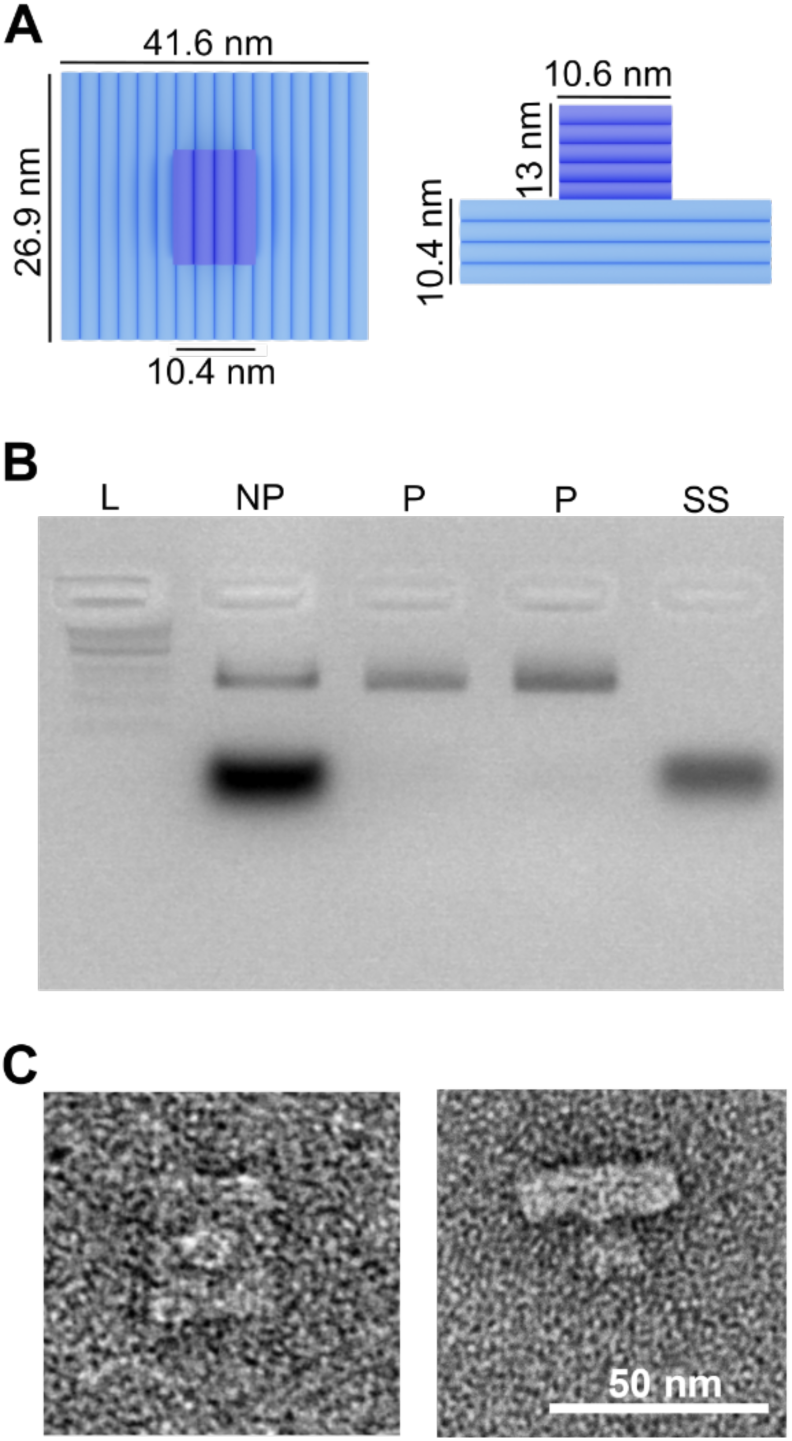
Structural design, assembly, and structural characterization of the DNA nanoprobe without lipid anchors. (A) Top and side view of the T-shaped DNP with nominal dimensions. (B) Gel electrophoretic analysis of non-purified DNP (NP), purified product (P), and excess staple strands (SS), using staining with ethidium bromide. (C) Transmission electron micrographs of negatively stained DNP displaying both its top (left) and side view (right).

The DNA nanostructure version lacking cholesterol tags was self-assembled by the origami method via the programmable folding of a long single-stranded DNA scaffold and shorter oligonucleotide staple strands that sequence-specifically hybridize to the scaffold.^72^ The staple sequences are provided in Tables S2 and S3, and the 2D DNA map and CanDo structure simulations in Figures S4 and S5, while the annealing protocol is in Table S4. Successful assembly as the DNA nanostructure was confirmed by agarose gel electrophoresis where the defined gel band for the product migrated higher than the scaffold (Figure 2B, Figure S6). The self-assembly product was purified from excess staple strands by cutting out the gel band and eluting the DNA structure (Figure 2B) for further structural analysis.^73^

The dimensions of purified DNP were determined with transmission electron microscopy. Analysis of the negatively stained samples (Figure 2C, Figure S7) revealed that the baseplate was 32.1 ± 0.5 nm in length, 41.2 ± 0.5 nm in width, and 14.5 ± 0.6 nm in height (n = 29). The width is in very good agreement with the expected value of 41.6 nm. The length is higher than the nominal dimension of 26.9 nm likely due to the extra nucleotide scaffold loops which can increase the duplex length. The observed higher height is a perspective effect due to the slight torsion of the DNP baseplate relative to the tip (Figure S5). By comparison, the experimentally determined tip dimensions were 9.9 ± 0.5 nm in width, 10.8 ± 1.1 nm in length, and 13.0 ± 1.0 nm in height (n = 20) which are close to the nominal values.

### Fabrication of DNA nanostructures with cholesterol membrane anchors

We fabricated than 20 anchor-modified DNP variants to explore how the steric accessibility of cholesterols influences binding to vesicles. The lipidated DNA nanostructures were self-assembled via origami in the presence of the cholesterol-modified DNA oligonucleotides (see Methods, Table S3). These strands hybridized to complementary DNA handles at designed locations of the nanostructure (Table S2) to position the cholesterol tags close to the DNP surface (Figure S1). The folding yield of individual DNPs was determined by gel electrophoresis (Figure S8) and ranged in relative terms from 100% (no-cholesterol DNP) to 31 ± 1.8 % (Figure S9), as determined by quantifying the intensity of the gel bands. Group DNP-4 had the lower yields likely due to the formation of dimers known to occur when cholesterols are on planar non-recessed DNA surfaces.^44, 49^

### Unilamellar vesicles

Small and large unilamellar vesicles composed of POPC and DOPC/DOPE/Chol (Figure 1C, Table S1) were generated via extrusion through filters with 200 nm, or 30 nm pores followed by tip sonication (see Methods). The vesicle populations’ size distribution and particle concentration were determined with dynamic light scattering (DLS). The analysis confirmed distinct and largely non-overlapping distributions for LUVs and SUVs of both membrane compositions (Figure 3, Figure S10). LUVs and SUVs of POPC composition had mean diameters of 192 ± 2 nm and 34 ± 4 nm. Similarly, LUVs and SUVs of DOPC/DOPE/Chol had mean diameters at 187 ± 2 nm and 58 ± 4 nm, respectively. The latter average value is higher than expected, likely due to the tendency of DOPC/DOPE/Chol membranes to fuse after vesicle extrusion.^74, 75^

**Figure 3.**
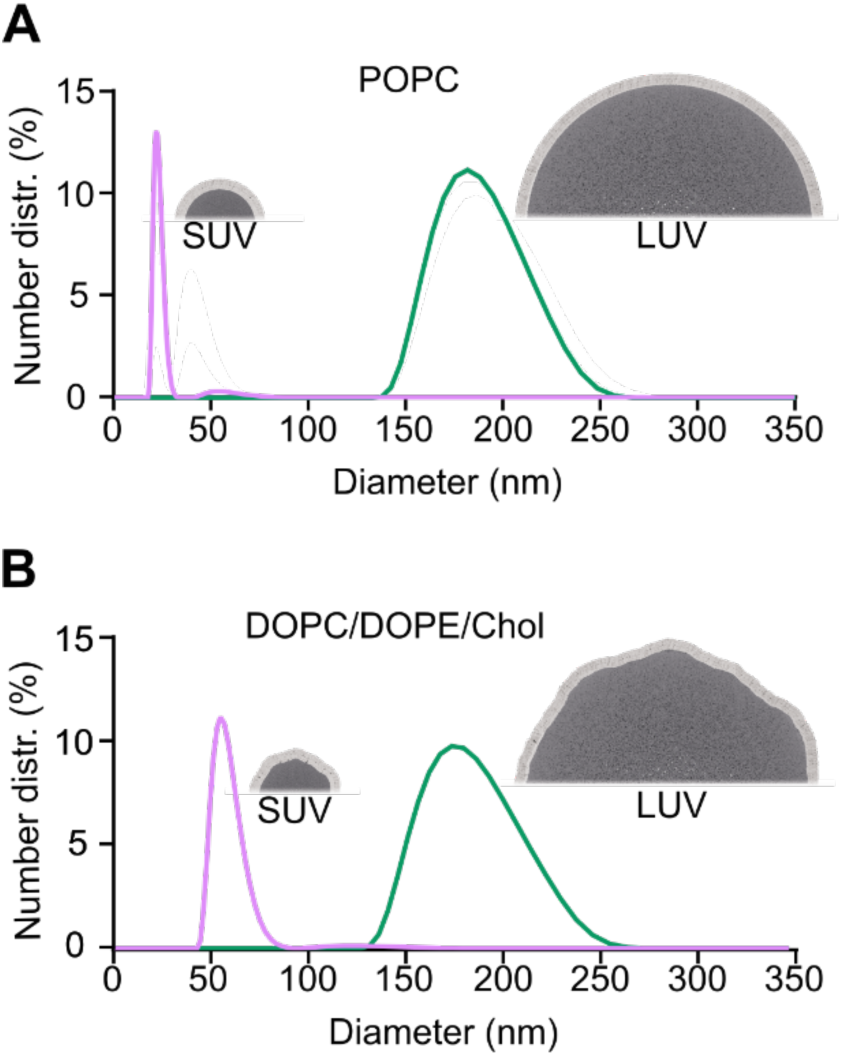
Hydrodynamic diameter distribution of the vesicle populations as determined with dynamic light scattering. (A) SUVs and LUVs with POPC membrane have means of 22 nm and 182 nm, respectively. (B) SUVs and LUVs composed of DOPC/DOPE/Chol (molar ratio of 2:1:2) have means of 51.2 nm and 183 nm, respectively. Additional DLS data are shown in Figure S10.

### Screening of DNP-vesicle binding using a gel-shift read-out

To understand how steric effects influence nanostructure-membrane interaction, we screened a total of 80 different combinations of parameters resulting from 20 lipidated DNA nanostructures, and four vesicle types of different curvature. For the assay, DNPs were incubated for 1 h with SUVs at a molar ratio of 1:1, and with LUVs at a ratio of 44:1 to maintain a ratio of one origami per approximately 2800 nm^2^ of outer leaflet surface. A low-magnesium buffer minimized nonspecific ionic binding of the vesicles with zwitterionic lipid headgroups.^52^ The vesicle-incubated samples and buffer-incubated DNP controls were screened via electrophoretic gel shift to determine the extent of vesicle. The results for POPC vesicles and the four groups DNP-1 to DNP-4 are summarized in Figure 4A-D. The gel shift analysis discriminated free and fast migrating DNA nanostructures from larger, vesicle-bound DNA nanostructures migrating slower in an upshifted gel band. This is exemplarily illustrated for nanostructure DNP-1.4 where DNP bands for SUVs and LUVs are upshifted compared to non-incubated DNP (Figure 4A, DNP-1.4, three lanes at right); the vesicle-induced gel shift depended on the vesicle size. Negative control DNP-0 without cholesterol did not show any gel upshift (Figure 4A, DNP-0, three leftmost lanes). To quantify the binding extent, scanned gel band intensities for each vesicle-bound DNP were normalized to the DNA nanostructure which had been incubated with vesicle-free buffer (see Methods). The binding extents are listed in Table S5 and summarized for POPC vesicles as line plots for groups DNP-1 to DNP-4 (Figure 4A-D).

**Figure 4.**
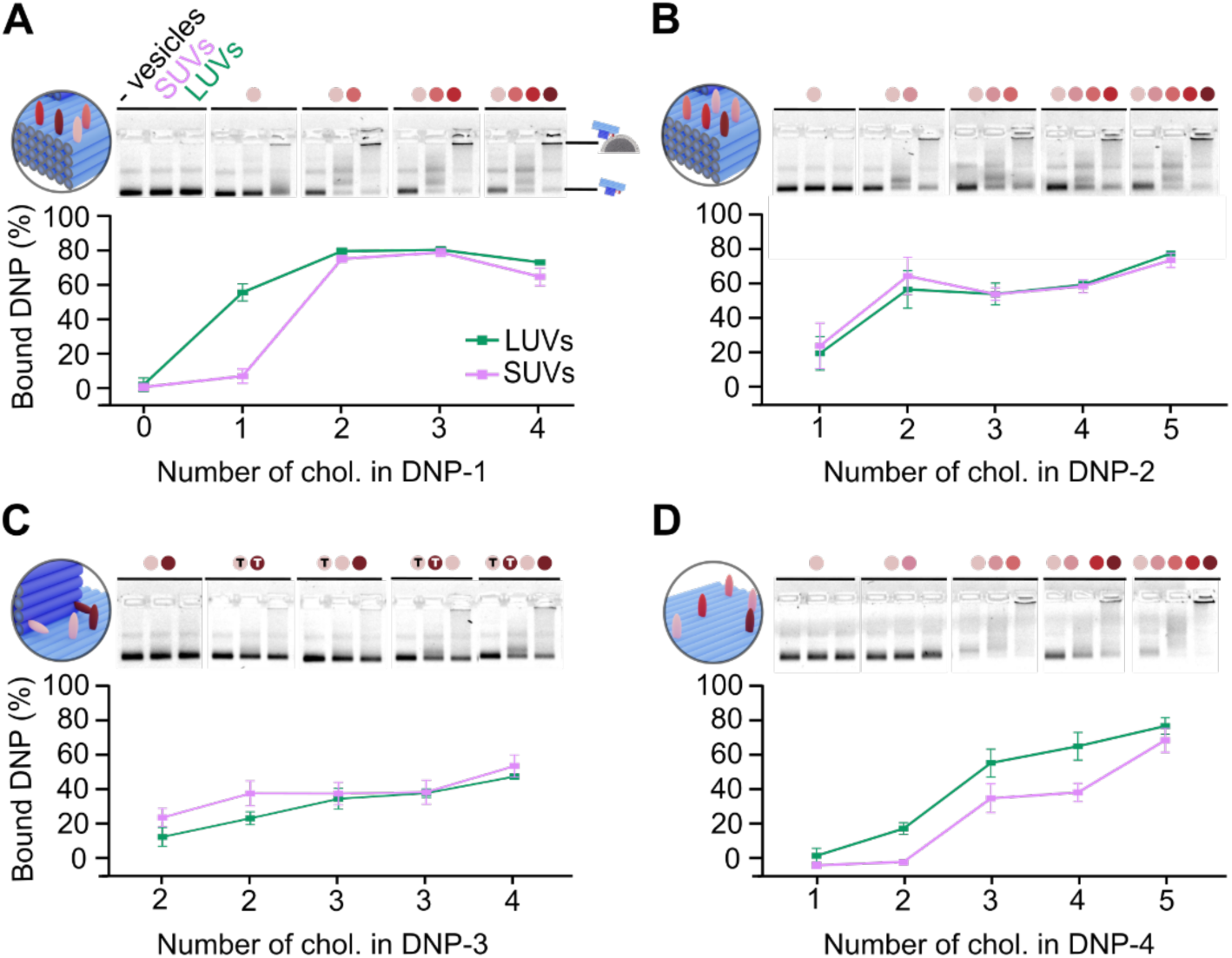
Binding of lipidated DNPs to POPC bilayer vesicles in dependence of cholesterol number and position, and vesicle size. Agarose gel electrophoresis and quantitative analysis on vesicles binding for nanostructure groups of (A) DNP-1, (B) DNP-2, (C) DNP-3, and (D) DNP-4. The agarose gels show each three lanes for free DNP, DNP after SUV incubation, and DNPs after LUV incubation, as visually linked by a horizontal line. The number and position of cholesterols in DNPs is indicated by red-colored dots at the top of each three lanes. The color-coded cholesterols in DNP are schematically illustrated to the left of each gel in the panel. In C, the “T” on colored dots represents the location of cholesterol at the DNP’s tip. The data points represent averages and the standard error from at least three independent experiments.

Comparing the line plots for DNP binding to POPC vesicles (Figure 4) reveals several key insights. (i) Higher binding yields are quantified to correlate with higher cholesterol number, in line with expectations, while the binding degree for a given cholesterol number strongly depends on their position, as illustrated for two cases. For DNP-1 nanostructures, two cholesterols yield over 79.1 ± 1.8 % binding to LUVs (Figure 4A) while the same cholesterol number in DNP-3 and DNP-4 results in low binding at 13.8 ± 6.5 and 17.1 ± 3.4 %, respectively (Figures 4C,D; Table S5, DNP-1.2, DNP-3.2, DNP-4.2). This difference can be partly explained by diverging steric accessibilities, such as for corner-positioned DNP-1 vs. tip-proximal DNP-3 (Figures 4A, C). However, in the case of DNP-4, the highly accessible planar underside of the DNP yields lower binding (Figure 4D) than equally or less accessible corner-to-tip proximal cholesterols of DNP-3 and DNP-4 (Figure 4A,B, 2 cholesterols). Preferred binding to cholesterols at the corners as opposed to the center has been reported for flat DNA nanostructures.^49, 54^ Our findings on preferred binding to the corner is surprising as the tip of the DNP structure was anticipated to sterically block access and binding of the vesicles.

In a further remarkable finding (ii), DNA nanostructure DNP-1.1 size-discriminates vesicles by efficiently binding to large LUVs but poorly to SUVs (Figure 4A) with an 8.2 ± 0.2 -fold difference (Table S5) and an apparent *K*_d_ for LUV interaction of 5.4 ± 1.2 x 10^-5^ M (Figure S11). This discrimination results from the cholesterol’s edge-corner-location as well as proximity to the DNP tip, as indicated by two controls. In the first, a single cholesterol located at the edge-corner but at the underside of the tip-free baseplate (Figure S1, S2) does not yield LUV binding (Figure S12). In the second control, a single cholesterol at a corner opposite to the initial cholesterol site (Figure S1) restores high LUV binding and effective discrimination against SUVs (Figure S12). The binding specificity is not a kinetic effect, as shown by the same binding extents following 15 h incubation (Figure S13). In contrast to successful discrimination, no binding difference is observed for group DNP-1 nanostructures with two or more cholesterols (Figure 4A) as well as group DNP-2 and DNP-3 nanostructures (Figure 4B,C).

Both findings of (i) poor vesicle binding to planar DNA structures, and (ii) efficient LUVs binding to DNA nanostructures with a nearby DNA tip can be reconciled by a unifying model. In the model, a single anchor is not sufficient for vesicle binding, likely as the entropic cost of forming the binary complex is not overcome. However, the presence of the DNP tip provides an additional interface bilayer binding (Figure S2) to stabilize the complex, as confirmed by the DNP control data (Figure S12). In other support, LUV vesicles with a larger and more deformable membrane bind the DNP-1.1 structure more than SUVs with a smaller and stiffer surface membrane area (Figure S2, Figure 4A). With two or more membrane anchors, binding is strong enough also for SUVs (Figure 4A). Our model reflects the complex interplay of factors also found for DNA structures that electrostatically bind to planar membranes,^76^ while it contrasts to poor protein binding to larger vesicles due to the different biomolecular geometries.^77^

Analyzing binding of DNP to DOPC/DOPE/Chol vesicles (Figure S14) confirmed results found for POPC vesicles, such as (i) a higher binding efficiency for higher cholesterol numbers, and the major role of cholesterol positions in binding. However, the separation of SUV and LUV binding previously observed for (ii) DNP-1.1 was less pronounced for DOPC/DOPE/Chol bilayers (compare Figure 4A with Figure S14A). This may be due to the smaller size difference between SUVs and LUVs of DOPC/DOPE/Chol than POPC composition (Figure 3). Another reason is that the mixed composition DOPC/DOPE/Chol bilayers allow SUVs and LUVs to exhibit heterogeneous and localized curvatures, thereby leveling the effects of global curvature.^64^ This latter argument is supported by the higher binding of DOPC/DOPE/Chol LUVs compared to POPC LUVs across all lipidated DNA nanostructure groups (Figure S15). The considerable role of localized membrane curvatures and lipid composition for 3D nanostructure binding constitutes finding (iii).

### DNP-vesicle interaction visualized with high-resolution fluorescence microscopy

As electrophoresis probes vesicle ensembles, we used a high-resolution single-molecule method to offer more detailed insight into how DNP binding extent depends on the actual vesicle diameter. As additional motivation, we sought to define the vesicle size threshold for selective binding of POPC LUVs to DNA nanoprobe DNP-1.1. As the method of choice, we selected fluorescence-based direct stochastic optical reconstruction microscopy (dSTORM)^78^ given its capacity to produce high-quality high-resolution single-molecule data.^79^

We first applied dSTORM to characterize SUV and LUV populations of POPC vesicles. Vesicles carrying Cy5-labeled lipids and biotinylated-lipid were immobilized via biotin-neutravidin bonds onto glass slides carrying a thin film of biotinylated poly(ethylene glycol)(PEG) and non-tagged PEG which reduces non-specific binding. The dSTORM micrographs showcase the clear size differences between SUVs and LUVs (Figure 5A,B) whereby each image pixel represents a single active fluorophore (Figure 5, top-right insets). Quantifying the diameter of the pixel clusters yielded the distribution of SUVs and LUVs (Figure 5, bottom-left insets). The averages of 57.3 ± 0.3 nm for SUVs and 170.3 ± 1.1 nm for LUVs. This is smaller than the NTA-derived averages of 75 ± 2 nm and 225 ± 12, respectively (Figure S10), as vesicle binding to surface is biased towards smaller vesicles due to the lower drag forces and faster diffusion.^80, 81^ Indeed, LUVs for dSTORM analysis were extruded with 400 nm filters instead of 200 nm to achieve immobilization of larger vesicles.

**Figure 5.**
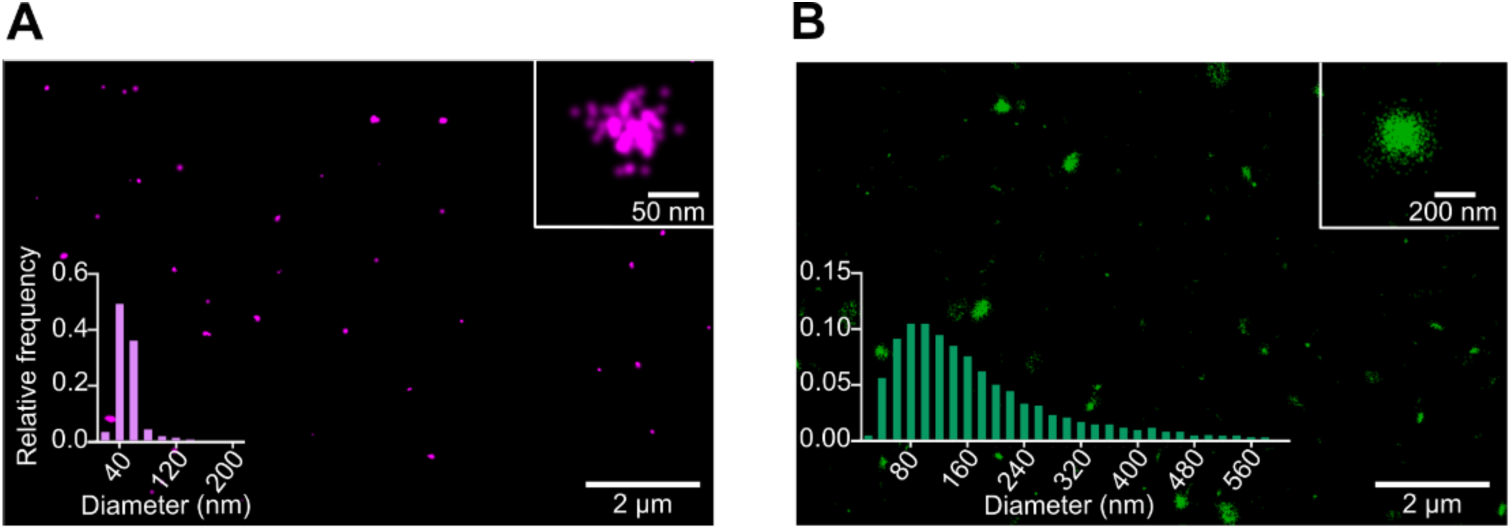
Single-molecule localization microscopy analysis of POPC vesicles. (A,B) dSTORM micrographs of Cy5-lipid and biotin-lipid-doped (A) SUVs and (B) LUVs immobilized via neutravidin-biotin bonds onto biotin-PEG/PEG-coated glass slides. Insets show single vesicles (top-right) and vesicle diameter distributions (bottom-left). LUVs and SUVs were prepared by extrusion with a filter of 400 nm or a filter of 30 nm followed by tip sonication, respectively. The diameters were obtained from the radius of gyration of the clustered localizations.

After visualizing vesicles, we explored size-selective binding of POPC large vesicles to DNA nanoprobe DNP-1.1 with single-particle resolution. dSTORM micrographs confirmed the selective binding of ATTO488-labeled DNPs-1.1 to LUVs but not SUVs (Figure 6C, Figure S16). Furthermore, positive control DNP-1.3 bound both vesicle types (Figure 6B, Figure S16) while negative control DNP-0 only showed negligible colocalization with either vesicle (Figure 6A, Figure S16). To quantify size-selective interaction, micrograph results were plotted as scatter plots (Figure 6D,E) whereby each dot reports on the extent of DNP binding as a function of vesicle diameter (Figure 6D,E). According to this analysis, the binding of positive control DNP-1.3 was similar across a wide range of vesicle diameters for SUVs and LUVs (Figure 6D). However, DNP-1.1 binding was poor on SUVs while increasing with the diameter of the LUVs with a threshold for binding at around 40 nm (Figure 6E). The binding extent for the negative control at about 5% (Figure S17). These findings further support the specificity of the cholesterol-modified DNA nanostructures in their interactions with vesicles.

**Figure 6.**
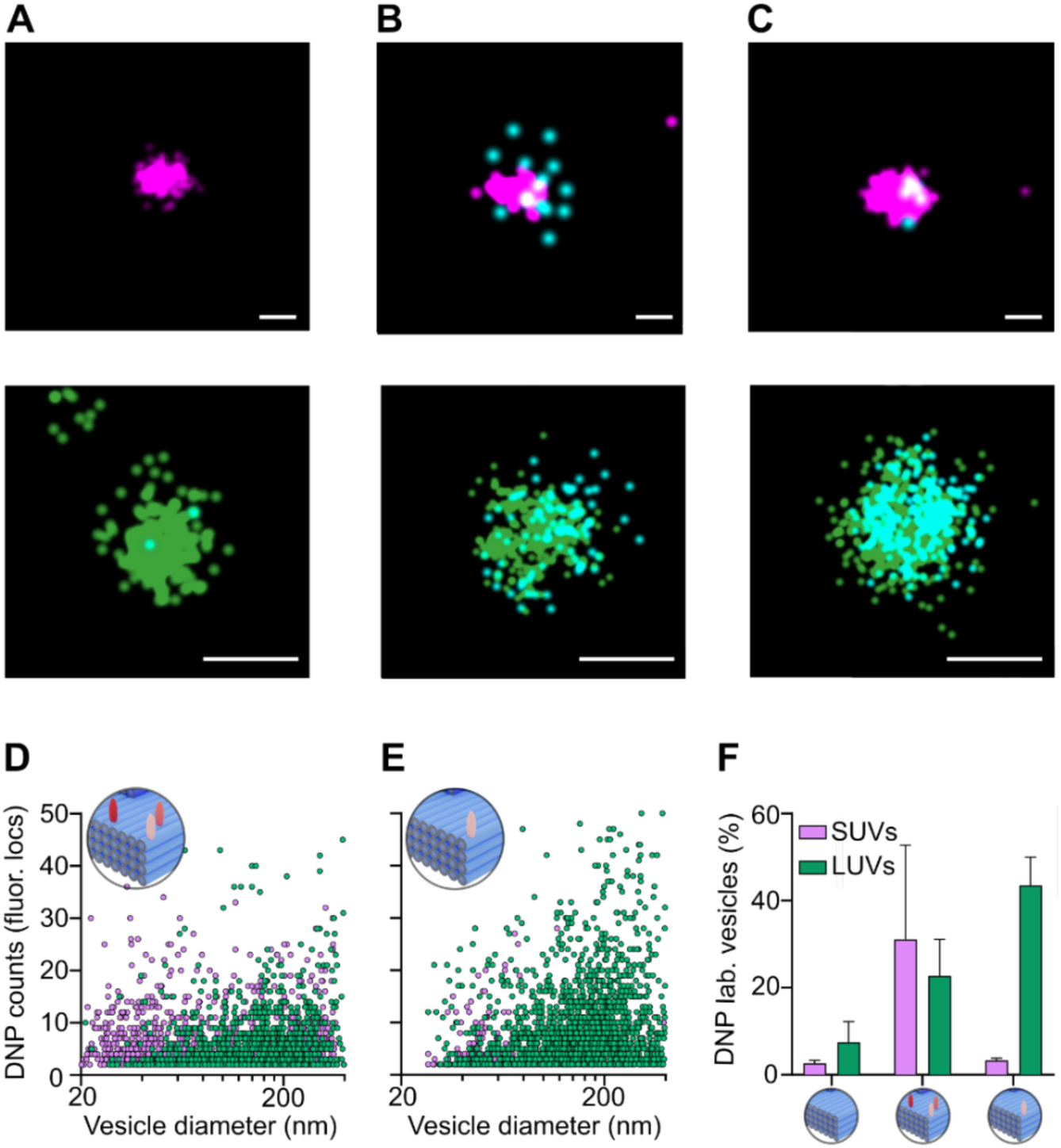
dSTORM analysis confirms the size-selective binding of POPC membrane vesicles to DNA nanoprobe DNP-1.1. (A,B,C) dSTORM micrographs of isolated SUVs (top panels) and LUVs (bottom panels) incubated with (A) DNP-0 without lipid anchor, (B) DNP-1.3 carrying three cholesterol anchors at the corner of the baseplate, and (C) DNP-1.1 with one cholesterol at the corner. DNPs and vesicles were labeled with fluorophores ATTO488 and Cy5, respectively. The fluorophore signals for SUVs are color-coded in magenta, LUVs in green, and DNPs in light blue. The scale bars are 50 nm for SUVs and 200 nm for LUVs. (D,E) Scatter plots for the dSTORM analysis of (D) DNP-1.3 and (E) DNP-1.1 binding which correspond to panels B and C, respectively. Each dot in a scatter plot represents the count of fluorescence-based DNP localizations within a vesicle (vertical axis) as a function of the corresponding vesicle diameter (horizontal axis). Signals from SUVs and LUVs are represented as magenta and green, respectively. (F) Bar plots of accumulated DNP signals for negative control DNP-0, DNP-3.1, and DNP-1.1 (left to right) for SUVs and LUVs within a radius of 300 nm around its center of gyration. The data for each condition represent the average and the standard error from at least three fields of view.

## CONCLUSIONS

Reflecting the technologically powerful role of biomimetic DNA nanostructures in synthetic biology and research, our study has determined how steric interactions underpin the formation of these hybrid DNA-lipid nanostructures. By exploring a wide parameters space using read-out including super-resolution microscopy, our study has yielded three main insights. First, we elucidated how DNA nanostructure-membrane interactions are influenced by the number and the geometric position of the membrane anchors in 3D geometries as opposed to flat nanostructures.^32, 33, 49, 50, 54^ To maximize insight, the membrane anchors were placed close to the DNA nanostructure surface to force tight interaction to lipid bilayer vesicles.^50^ This contrasts with previous studies where membrane anchors have been attached via polymeric and DNA linkers of up to 10 nanometers in length on flat^32, 33, 49, 50, 54^ and curved^19, 23, 28^ DNA nanostructures. While long linkers increase conformational flexibility and hence improve membrane interaction, they can blur the picture of how anchor position influences binding^19, 20, 23, 33, 49, 50^ even though molecular accessibility of lipid anchors controls vesicles binding^54^ and the related phenomenon of DNA duplex aggregation.^44^ However, 3D geometries of DNA nanostructure are relevant parameters for biological applications such as shown by in vivo uptake of DNA origami into human cells via the pathway of small endosome vesicles for drug delivery applications.^31^ Our findings therefore provide key information which help rationalize nanostructure-bilayer interactions.

Secondly, we have provided experimental evidence of how anchor-mediated DNA nanostructure-lipid bilayer interaction is governed not solely by membrane anchors. We propose that, an important, but usually disregarded interactions between non-modified nanostructure surfaces and other membrane segments. These non-anchored interactions likely act cooperatively with the membrane anchors stabilizing the entire interaction. The model is supported by several of our experimental findings, the most prominent being the size-selective binding of large unilamellar vesicles to a DNA nanostructure with a size cutoff above 40 nm vesicle diameter. This nanostructure-based size discrimination between vesicles may, with further development, be exploited to distinguish diagnostically relevant exosome membrane vesicles in the size range of 30 to 150 nm^67, 68^ and thereby fill a gap in the diagnostic tool set. As exosomes have distinct biological properties depending on their size, instrument-free separation is of considerable interest to complement currently used ultracentrifugation and size exclusion chromatography^65^ to allow tumor-specific exosomal biomarkers and exosome-based liquid biopsies.^82, 83^ Our DNA nanostructure-based principle to size-selectively bind vesicles may be able to address this demand. Indeed, other DNA nanostructures capable of measuring different vesicle curvatures have recently been developed for simple fluorescence read-out.^27^

The third insight is the major role of lipid composition and membrane curvature in anchor-mediated nanostructure binding. We reveal that vesicle size only matters for binding to homogeneous POPC membranes while for DOPC/DOPE/Chol the global curvature is dominated by fine-grained localized bilayer curvatures. The role of global and fine-grained previous findings in dependence of lipid composition complements a previous finding on better DNA structure binding to membranes of higher cholesterol content^54, 84^ to better predict and tailor DNA-bilayer interactions. In conclusion, our study extends programmable assembly interaction within DNA nanotechnology towards lipid membranes and lays the foundation for new nanostructures for applications in synthetic biology, biophysical research, and the diagnosis and purification of exosomes in complex mixtures.

## METHODS

### Materials

Unmodified, 3′-cholesterol-tetraethyleneglycol-modified, and 3′-ATTO488-modified DNA oligonucleotides were purchased from Integrated DNA Technologies (Belgium). 5′-cholesterol-tetraethyleneglycol-modified DNA oligonucleotides were purchased from Eurogentec (Belgium). The scaffold M13mp18 was procured from tilibit nanosystems (Germany). Lipids 1-palmitoyl-2-oleoyl-*sn*-glycero-3-phosphocholine (POPC), 1,2-dioleoyl-*sn*-glycero-3-phosphocholine (DOPC), 1,2-dioleoyl-*sn*-glycero-3-phosphoethanolamine (DOPE), cholesterol (Chol), 1,2-dioleoyl-*sn*-glycero-3-phosphoethanolamine-N-(Cyanine5) (Cy5-DOPE), and 1,2-dioleoyl-*sn*-glycero-3-phosphoethanolamine-N-(cap biotinyl) (biotin-DOPE) (structures in Table S5) were purchased from Sigma-Aldrich (UK). Agarose powder was purchased from Invitrogen (UK). Purple ‘no SDS’ loading dye was purchased from New England Biolabs (NEB). PEG:bioPEG microscopy chips were purchased from Oxford Nanoimaging (Oxford, UK). All other solvents and reagents were purchased from Sigma-Aldrich.

### Design and folding of the DNA nanoprobe

The DNA nanostructures were designed in Cadnano2 using the square lattice mode.^69^ To avoid twists resulting from the underwinding of the DNA helices in the square lattice arrangement, a base pair deleted every 56 nucleotides in all duplexes. This twist correction and the rigidity of the structure were verified with CanDo.^81^ The design furthermore included loops formed by sixteen unhybridized nucleotides of the scaffold strand at the duplex ends to prevent commonly occurring aggregation of DNA nanostructures. The design also ensured that cholesterol moieties of the modified strands were positioned close to the DNA baseplate or tip surface as illustrated in Figure S6. For dSTORM analysis, eight ATTO488 fluorophores were incorporated into the DNP by hybridizing the fluorophore-modified oligonucleotides to the edges of the baseplate.

DNPs were self-assembled by mixing M13mp18 scaffold (final concentration of 20 nM) with staple strands in 10-fold excess and optionally 5′-cholesterol modified DNA oligonucleotides in 50-fold excess, or 3′-cholesterol-modified oligonucleotides in 100-fold excess over the scaffold concentration within TAE buffer containing 16 mM MgCl_2_. The sequences of the non-modified staple strands are provided in Table S2 and those of chemically modified strands in Table S3. For microscopy experiments, the assembly mix included ATTO488-labeled DNA oligonucleotides in 200-fold excess. To achieve self-assembly, the mixture was subjected to thermal annealing from 95 °C to 4 °C in a thermocycler using an annealing program detailed in Table S4. Following folding, excess staple strands were removed via spin filtration with an Amicon® Ultra, MWCO 50 KDa filter (Millipore) using 3 x centrifugation at 7000 g for 4 min each and exchanging the folding buffer for 1 × TAE supplemented with 3 mM MgCl_2_ and 300 mM NaCl. After purification, DNP concentrations were determined by measuring the absorbance at 260 nm with a DS-11 spectrophotometer (DeNovix, USA) assuming a calculated extinction coefficient of 8.96 × 10^7^ M^−1^ cm^−1^.

### Transmission Electron Microscopy

The DNP structure was subjected to TEM analysis to confirm shape and dimensions. The TEM samples were prepared by adding cholesterol-free DNP solution (5 nM, 6 μL) for 10 sec onto glow discharged Cu300 mesh grids coated with carbon. The samples were then subjected to negative stain with a 2% uranyl acetate solution. TEM imaging was performed using a JEM-2100 electron microscope (JEOL, Japan) operating at 200 kV. Images were captured using an Orius SC200 camera.

### Vesicle preparation

Small and large unilamellar vesicles (SUVs and LUVs) were prepared to study the interaction between DNP and vesicles using an electrophoretic gel shift assay. The preparation of SUVs and LUVs followed a sonication/extrusion protocol using POPC and DOPC/DOPE/Chol (2:1:2). Lipids (total of 9 mg) were dissolved in chloroform, followed by drying under nitrogen gas to form a thin lipid film which was then subjected to vacuum for at least 3 h to remove any traces of chloroform. The lipid film was then resuspended in 1 mL of 1 × TAE supplemented with 300 mM NaCl. The solution was shaken on a thermomixer (Eppendorf Ltd, UK) at 800 rpm for 30 min at 30°C. To obtain LUVs, the vesicle suspension was extruded 31 times through a Whatman Nucleopore™ track-etched polycarbonate membrane (Merck, Germany) with 200 nm pore size using a Mini-Extruder (Avanti Polar Lipids, Inc., USA). To produce SUVs, 500 μL of the LUVs solution was diluted with 500 μL of 1 × TAE supplemented with 300 mM NaCl and extruded 31 x through a 50 nm pore size membrane, and 15 x through a 30 nm membrane. The SUVs were left in ice and subjected to tip sonication with a Sonifier® 150D (Branson Ultrasonics, USA) at 60 w, 22 ± 1.3 kHz, with 10-seconds on-off pulses for a total duration of 10 minutes. Following sonication, the SUVs were spin filtered through an MWCO 300 KDa filter (Vivaspin®, Sartorius AG, Germany) at 7000 g for 4 min to remove any contamination from the sonication tip, and the resulting filtrate was collected. Vesicles were kept at 4 °C for up to one week and were characterized by DLS and NTA before use.

For dSTORM experiments, vesicles were prepared by suspending the lipid film in 1 x TAE with 572 mM NaCl to match the osmolarity of the buffer used during dSTORM. To facilitate immobilization and fluorescence-based visualization, membrane lipids included a molar fraction of 0.5% biotin-DOPE, and 0.3% of Cy5-DOPE for SUVs and 0.03% for LUVs. LUVs for microscopy experiments were prepared by extruding the lipid solution 31 times using a 400 nm pore-size membrane. After extrusion, the LUVs were subjected to three rounds of spin filtration using an MWCO 1000 kDa filter at 12,000 g for 2 min. This filtration step helped reduce the population of sub-200 nm vesicles produced during extrusion.

### Dynamic Light Scattering

DLS analysis was conducted to determine the size distribution and concentration of unlabeled vesicle samples. The measurements were performed on a Zetasizer Ultra (Malvern Instrument, Malvern, UK) operating with a 4 mW HeNe laser at a wavelength of 633 nm. The size distribution of the vesicles was measured using non-invasive Back Scatter (NIBS) with a scattering angle of 173 degrees. A total of 30 measurements were taken at a temperature of 25 °C, and the results were averaged. Particle concentrations were determined by multi-angle dynamic light scattering (MADLS)^85^ by measuring at detection angles of 173, 13, and 90 degrees, 30 times each. The collected data were interpreted using the ZS XPLORER software provided by the manufacturer. The size distribution of the samples was determined using an L-curve-based fitting algorithm, considering the buffer viscosity as that of water at 25 °C.

### Nanoparticle Tracking Analysis

To determine the size distribution of Cy5-labeled vesicles, NTA analysis as DLS is incompatible as the laser beam used for scattering also excites the Cy5 fluorophores. NTA analysis was carried out using a NanoSight LM10 (NanoSight, Amesbury, United Kingdom) equipped with a 488 nm laser and operating in scattering mode. Before measurements, vesicle samples were diluted to approximately 10^8^ particles mL^-1^ to ensure reliable determination of the size distributions between different samples. For each sample, a total of five videos of 60 sec duration and frame rate of 30 Hz were recorded. The videos were subsequently analyzed using the built-in NTA 3.2 software considering buffer viscosity as that of water at the recorded experiment’s temperature.

### Agarose Gel Electrophoresis

Binding of DNPs to vesicles was assessed using electrophoresis with 2% agarose gel in 0.5 x TBE buffer supplemented with 100 mM NaCl buffer and 0.5 µg mL^-1^ ethidium bromide. Before loading onto the gel, DNPs were incubated at a concentration of 6.1 nM a ratio of 1:1 with SUVs or a ratio of 44:1 with LUVs to maintain an accessible lipid surface of 2827 nm^2^ of per DNP. The vesicle concentrations were adjusted based on their DLS-measured average diameter. After incubation, “no SDS” loading dye (New England Biolabs, UK; 4 µL) was added to each sample (15 µL, approximately 320 ng of DNA), and the mixture was loaded onto the gel. Scaffold M13mp18 strand (300 ng) was loaded onto each gel to compare DNP yields between different gels. The electrophoresis was conducted in an ice-water bath, at a voltage of 3 V per cm for 210 min using 0.5 × TBE buffer supplemented with 100 mM NaCl and 50 ng mL^-1^ of ethidium bromide to prevent gel destaining during electrophoresis. Following electrophoresis, gels were imaged under UV illumination (c Series Imaging Systems, Azure Biosystems, USA). Gel images were analyzed using ImageJ^86^ to quantify the DNP binding. Image background was subtracted using a rolling ball algorithm, and band intensities for each DNP-vesicle combination were quantified and normalized against the intensity of the DNP in the absence of vesicles. The folding yield was obtained by using the intensity of the scaffold band as reference.

### Preparation of microscopy slides

Microscopy chips with four microfluidic channels and glass surfaces coated with a PEG and biotin-PEG thin film (Oxford Nanoimaging, UK) were washed three times with TAE buffer and subsequently incubated with a solution of neutravidin (1 mg/mL, 20 μL, PBS) for 15 min. After gently washing the microfluidic channels with TAE buffer, surfaces were passivated by incubation with a casein solution (1 mg mL^-1^, 20 μL, PBS) for 30 min. The channels were gently rinsed with TAE buffer, followed by a suspension of SUVs (vesicle concentration of 0.75 nM, 30 μL) or LUVs (0.2 nM, 30 μL). After 1 min of incubation, unbound vesicles were removed by washing with TAE buffer supplemented with 572 mM NaCl. To test binding of DNP to vesicles, SUVs, and LUVs were incubated for 1 h with ATTO488-labeled DNP variants at a concentration of 2 nM, followed by washing with TAE buffer containing 572 mM NaCl. All buffers and components used for slide preparation and microscopy were filtered with a 0.2 uM polyethersulfone syringe filter.

### Direct Stochastic Optical Reconstruction Microscopy

Lipid vesicles were imaged with dSTORM to determine their size distribution and to quantify the binding of DNPs to vesicles. dSTORM was conducted using a Nanoimager S Mark II microscope from ONI (Oxford Nanoimaging, Oxford, UK) equipped with a 100×, 1.4NA oil immersion objective, an XYZ closed-loop piezo 736 stage, and dual emission channels split at 640 nm. Samples were imaged in buffer MEA/GLOX (10 mM mercaptoethanolamine, 5.6 mg mL^-1^ glucose oxidase, 340 μg mL^-1^ catalase, 10% (w/v) glucose).^87^ Images were acquired at 50 Hz, with an illumination angle of 53 degrees to image in total internal reflection mode. Imaging was carried out through sequential acquisition, with the first 13,000 frames recorded under illumination with 640 nm laser and then 10,000 frames under 473 nm laser illumination (2.5 kW cm^−2^). The power of the 640 nm laser was modulated between the imaged vesicles to adjust to the Cy5 fluorescence density of the different sizes of vesicles. Hence, the field of view for LUVs was initially bleached for 10 sec with a power of 4 kW and then imaged with a power of 2 kW cm^−2^ while for SUVs, the field of view was directly imaged with a power of 0.8 kW cm^−2^. The acquired images were processed using ONI’s Collaborative Discovery online analysis platform. All images were filtered using consistent parameters. Images were first corrected for drift through a redundant-cross-correlation algorithm.^88^ Then, localization was filtered based on parameters such as point spread function shape, photon count (>200 photons), and localization precision (< 30 nm) to minimize visual artifacts and remove low-precision localizations. The filtered Cy5 localizations were then clustered, and the radius of gyration of each cluster was calculated. Additionally, the number of ATTO488 localizations was counted in a radius of 300 nm from the center of gyration of each cluster.

## ASSOCIATED CONTENT

### Supporting Information

The Supporting Information is available free of charge on the ACS Publications website at DOI: 10.1021/acsnano.xxx. DNA sequences, combinations of oligonucleotides used to generate each construct, 2D maps, and 3D renderings, as well as additional experimental data on formation and characterization of DNA nanostructures and lipid vesicles, gel electrophoretic assay on the binding of DNA nanostructures to lipid vesicles, and single molecule localization microscopy.

## AUTHOR INFORMATION

### Author Contributions

Y.Z. designed the DNA nanostructure and conducted the TEM analysis. E.G. and J.C.G. devised and carried out all other experiments. J.C.G. and E.G. wrote a draft of the manuscript and S.H. edited the manuscript with input from E.G. and Y.X. E.G produced the manuscript figures. S.H. supervised the project.

## ACKNOWLEDGMENT

We thank Miguel Paez Perez for his contributions at an early stage of the project and valuable discussions.

## ABBREVIATIONS

DLS: Dynamic Light Scattering
DNP: DNA Nanoprobe
DOPC: 2-dioleoyl-sn-glycero-3-phosphocholine
DOPE: 1,2-dioleoyl-sn-glycero-3-phosphoethanolamine
dSTORM: direct stochastic optical reconstruction microscopy
LUV: Large Unilamellar vesicle
NTA: Nanoparticle Tracking Analysis
POPC: 1-palmitoyl-2-oleoyl-glycero-3-phosphocholine
SUV: Small unilamellar vesicle
TEM: transmission electron microscopy.

## ToC Graphic

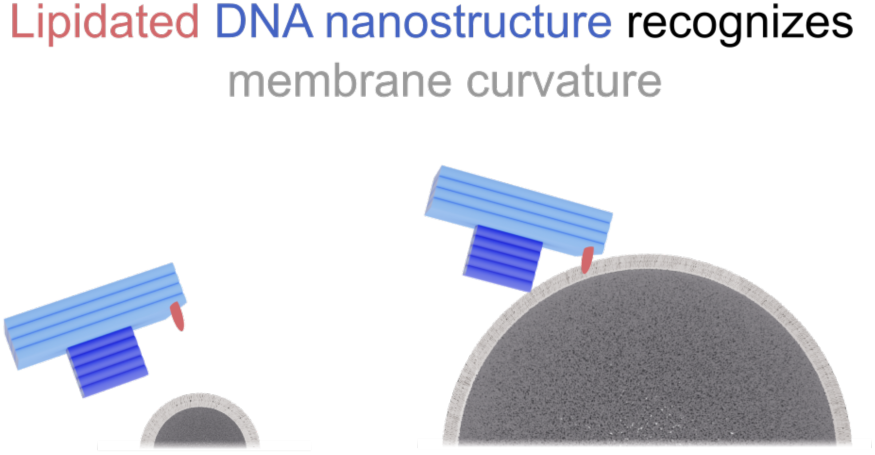

## REFERENCES

(1) Rothemund, P. W. K. Folding DNA to Create Nanoscale Shapes and Patterns. Nature 2006, 440, 297–302.

(2) Seeman, N. C.; Sleiman, H. F. DNA Nanotechnology. Nat. Rev. Mater. 2017, 3, 1–23.

(3) Fu, J.; Yang, Y. R.; Johnson-Buck, A.; Liu, M.; Liu, Y.; Walter, N. G.; Woodbury, N. W.; Yan, H. Multi-Enzyme Complexes on DNA Scaffolds Capable of Substrate Channelling with an Artificial Swinging Arm. Nat. Nanotechnol. 2014, 9, 531–536.

(4) Funke, J. J.; Dietz, H. Placing Molecules with Bohr Radius Resolution Using DNA Origami. Nat. Nanotechnol. 2016, 11, 47–52.

(5) Madsen, M.; Gothelf, K. V. Chemistries for DNA Nanotechnology. Chem. Rev. 2019, 119, 6384–6458.

(6) Douglas, S. M.; Dietz, H.; Liedl, T.; Högberg, B.; Graf, F.; Shih, W. M. Self-Assembly of DNA into Nanoscale Three-Dimensional Shapes. Nature 2009, 459, 414–418.

(7) Dietz, H.; Douglas, S. M.; Shih, W. M. Folding DNA into Twisted and Curved Nanoscale Shapes. Science 2009, 325, 725–730.

(8) Ke, Y.; Ong, L. L.; Shih, W. M.; Yin, P. Three-Dimensional Structures Self-Assembled from DNA Bricks. Science 2012, 338, 1177–1183.

(9) Benson, E.; Mohammed, A.; Gardell, J.; Masich, S.; Czeizler, E.; Orponen, P.; Högberg, B. DNA Rendering of Polyhedral Meshes at the Nanoscale. Nature 2015, 523, 441–444.

(10) Nummelin, S.; Kommeri, J.; Kostiainen, M. A.; Linko, V. Evolution of Structural DNA Nanotechnology. Adv. Mater. 2018, 30, 1703721.

(11) Hu, Y.; Niemeyer, C. M. From DNA Nanotechnology to Material Systems Engineering. Adv. Mater. 2019, 31, 1806294.

(12) Kuzyk, A.; Schreiber, R.; Fan, Z.; Pardatscher, G.; Roller, E. M.; Högele, A.; Simmel, F. C.; Govorov, A. O.; Liedl, T. DNA-Based Self-Assembly of Chiral Plasmonic Nanostructures with Tailored Optical Response. Nature 2012, 483, 311–314.

(13) Tikhomirov, G.; Petersen, P.; Qian, L. Fractal Assembly of Micrometre-Scale DNA Origami Arrays with Arbitrary Patterns. Nature 2017, 552, 67–71.

(14) Ong, L. L.; Hanikel, N.; Yaghi, O. K.; Grun, C.; Strauss, M. T.; Bron, P.; Lai-Kee-Him, J.; Schueder, F.; Wang, B.; Wang, P.; Kishi, J. Y.; Myhrvold, C.; Zhu, A.; Jungmann, R.; Bellot, G.; Ke, Y.; Yin, P. Programmable Self-Assembly of Three-Dimensional Nanostructures from 10,000 Unique Components. Nature 2017, 552, 72–77.

(15) Sontakke, V. A.; Yokobayashi, Y. Programmable Macroscopic Self-Assembly of DNA-Decorated Hydrogels. J. Am. Chem. Soc. 2022, 144, 2149–2155.

(16) Olzmann, J. A.; Carvalho, P. Dynamics and Functions of Lipid Droplets. Nat. Rev. Mol. Cell Biol. 2019, 20, 137–155.

(17) Altamura, E.; Milano, F.; Tangorra, R. R.; Trotta, M.; Omar, O. H.; Stano, P.; Mavelli, F. Highly Oriented Photosynthetic Reaction Centers Generate a Proton Gradient in Synthetic Protocells. PNAS 2017, 114, 3837–3842.

(18) Bridges, A. A.; Zhang, H.; Mehta, S. B.; Occhipinti, P.; Tani, T.; Gladfelter, A. S. Septin Assemblies Form by Diffusion-Driven Annealing on Membranes. PNAS 2014, 111, 2146–2151.

(19) Franquelim, H. G.; Khmelinskaia, A.; Sobczak, J.-P.; Dietz, H.; Schwille, P. Membrane Sculpting by Curved DNA Origami Scaffolds. Nat. Commun. 2018, 9, 811.

(20) Jahnke, K.; Huth, V.; Mersdorf, U.; Liu, N.; Göpfrich, K. Bottom-Up Assembly of Synthetic Cells with a DNA Cytoskeleton. ACS Nano 2022, 16, 7233–7241.

(21) Birkholz, O.; Burns, J. R.; Richter, C. P.; Psathaki, O. E.; Howorka, S.; Piehler, J. Multi-Functional DNA Nanostructures That Puncture and Remodel Lipid Membranes into Hybrid Materials. Nat. Commun. 2018, 9, 1521.

(22) Ahmad, M.; Prensky, H.; Balestrieri, J.; ElNaggar, S.; Gomez-Simmonds, A.; Uhlemann, A. C.; Traxler, B.; Singh, A.; Lopatkin, A. J. Tradeoff between Lag Time and Growth Rate Drives the Plasmid Acquisition Cost. Nat. Commun. 2023, 14, 2343.

(23) Zhang, Z.; Yang, Y.; Pincet, F.; Llaguno, M. C.; Lin, C. Placing and Shaping Liposomes with Reconfigurable DNA Nanocages. Nat. Chem. 2017, 9, 653–659.

(24) Rubio-Sánchez, R.; Barker, S. E.; Walczak, M.; Cicuta, P.; Michele, L. Di. A Modular, Dynamic, DNA-Based Platform for Regulating Cargo Distribution and Transport between Lipid Domains. Nano Lett. 2021, 21, 2800–2808.

(25) Xing, H.; Zhu, Y.; Xu, D.; Wu, R.; Xing, X.; Li, L. S. DNA Tetrahedron–Mediated Triplex Molecular Switch for Extracellular pH Monitoring. Anal. Chim. Acta. 2023, 1265, 341336.

(26) Silvester, E.; Vollmer, B.; Pražák, V.; Vasishtan, D.; Machala, E. A.; Whittle, C.; Black, S.; Bath, J.; Turberfield, A. J.; Grünewald, K.; Baker, L. A. DNA Origami Signposts for Identifying Proteins on Cell Membranes by Electron Cryotomography. Cell 2021, 184, 1110– 1121.

(27) Büber, E.; Schröder, T.; Scheckenbach, M.; Dass, M.; Franquelim, H. G.; Tinnefeld, P. DNA Origami Curvature Sensors for Nanoparticle and Vesicle Size Determination with Single-Molecule FRET Readout. ACS Nano 2023, 17, 3088–3097.

(28) Bian, X.; Zhang, Z.; Xiong, Q.; De Camilli, P.; Lin, C. A Programmable DNA-Origami Platform for Studying Lipid Transfer between Bilayers. Nat. Chem. Biol. 2019, 15, 830–837.

(29) Schneider, L.; Rabe, K. S.; Domínguez, C. M.; Niemeyer, C. M. Hapten-Decorated DNA Nanostructures Decipher the Antigen-Mediated Spatial Organization of Antibodies Involved in Mast Cell Activation. ACS Nano 2023, 17, 6719–6730.

(30) Huang, D.; Patel, K.; Perez-Garrido, S.; Marshall, J. F.; Palma, M. DNA Origami Nanoarrays for Multivalent Investigations of Cancer Cell Spreading with Nanoscale Spatial Resolution and Single-Molecule Control. ACS Nano 2019, 13, 728–736.

(31) Rajwar, A.; Shetty, S. R.; Vaswani, P.; Morya, V.; Barai, A.; Sen, S.; Sonawane, M.; Bhatia, D. Geometry of a DNA Nanostructure Influences Its Endocytosis: Cellular Study on 2D, 3D, and in Vivo Systems. ACS Nano 2022, 16, 10496–10508.

(32) Kocabey, S.; Kempter, S.; List, J.; Xing, Y.; Bae, W.; Schiffels, D.; Shih, W. M.; Simmel, F. C.; Liedl, T. Membrane-Assisted Growth of DNA Origami Nanostructure Arrays. ACS Nano 2015, *9*, 3530–3539.

(33) Kanwa, N.; Gavrilovic, S.; Brüggenthies, G. A.; Qutbuddin, Y.; Schwille, P. Inducing Lipid Domains in Membranes by Self-Assembly of DNA Origami. Adv. Mater. Interfaces 2023, 10, 2202500.

(34) Joesaar, A.; Yang, S.; Bögels, B.; van der Linden, A.; Pieters, P.; Kumar, B. V. V. S. P.; Dalchau, N.; Phillips, A.; Mann, S.; de Greef, T. F. A. DNA-Based Communication in Populations of Synthetic Protocells. Nat. Nanotechnol. 2019, 14, 369–378.

(35) Lyu, Y.; Wu, C.; Heinke, C.; Han, D.; Cai, R.; Teng, I.-T.; Liu, Y.; Liu, H.; Zhang, X.; Liu, Q.; Tan, W. Constructing Smart Protocells with Built-In DNA Computational Core to Eliminate Exogenous Challenge. J. Am. Chem. Soc. 2018, 140, 6912–6920.

(36) Samanta, A.; Hörner, M.; Liu, W.; Weber, W.; Walther, A. Signal-Processing and Adaptive Prototissue Formation in Metabolic DNA Protocells. Nat. Commun. 2022, 13, 3968.

(37) Arulkumaran, N.; Singer, M.; Howorka, S.; Burns, J. R. Creating Complex Protocells and Prototissues Using Simple DNA Building Blocks. Nat. Commun. 2023, 14, 1314.

(38) Chen, H.; Xu, W.; Shi, H.; Qiao, Y.; He, X.; Zheng, J.; Zhou, S.; Yang, X.; Wang, K.; Liu, J. DNA-Based Artificial Receptors as Transmembrane Signal Transduction Systems for Protocellular Communication. Angew. Chem. Int. Ed. 2023, 62, e202301559.

(39) Xing, Y.; Dorey, A.; Jayasinghe, L.; Howorka, S. Highly Shape- and Size-Tunable Membrane Nanopores Made with DNA. Nat. Nanotechnol. 2022, 17, 708–713.

(40) Xing, Y.; Rottensteiner, A.; Ciccone, J.; Howorka, S. Functional Nanopores Enabled with DNA. Angew. chem. Int. Ed. 2023, e202303103.

(41) Dey, S.; Dorey, A.; Abraham, L.; Xing, Y.; Zhang, I.; Zhang, F.; Howorka, S.; Yan, H. A Reversibly Gated Protein-Transporting Membrane Channel Made of DNA. Nat. Commun. 2022, 13, 2271.

(42) Zheng, H.; Li, H.; Li, M.; Zhai, T.; Xie, X.; Li, C.; Jing, X.; Liang, C.; Li, Q.; Zuo, X.; Li, J.; Fan, J.; Shen, J.; Peng, X.; Fan, C. A Membrane Tension-Responsive Mechanosensitive DNA Nanomachine. Angew. Chem. Int. Ed. 2023, e202305896.

(43) Ohmann, A.; Li, C.-Y.; Maffeo, C.; Nahas, K. Al; Baumann, K. N.; Göpfrich, K.; Yoo, J.; Keyser, U. F.; Aksimentiev, A. A Synthetic Enzyme Built from DNA Flips 107 Lipids per Second in Biological Membranes. Nat. Commun. 2018, 9, 2426.

(44) Ohmann, A.; Göpfrich, K.; Joshi, H.; Thompson, R. F.; Sobota, D.; Ranson, N. A.; Aksimentiev, A.; Keyser, U. F. Controlling Aggregation of Cholesterol-Modified DNA Nanostructures. Nucleic Acids Res. 2019, 47, 11441–11451.

(45) Jones, S. F.; Joshi, H.; Terry, S. J.; Burns, J. R.; Aksimentiev, A.; Eggert, U. S.; Howorka, S. Hydrophobic Interactions between DNA Duplexes and Synthetic and Biological Membranes. J. Am. Chem. Soc. 2021, 143, 8305–8313.

(46) Krishnan, S.; Ziegler, D.; Arnaut, V.; Martin, T. G.; Kapsner, K.; Henneberg, K.; Bausch, A. R.; Dietz, H.; Simmel, F. C. Molecular Transport through Large-Diameter DNA Nanopores. Nat. Commun. 2016, 7, 12787.

(47) Burns, J. R.; Göpfrich, K.; Wood, J. W.; Thacker, V. V.; Stulz, E.; Keyser, U. F.; Howorka, S. Lipid-Bilayer-Spanning DNA Nanopores with a Bifunctional Porphyrin Anchor. Angew. Chem. Int. Ed. 2013, 52, 12069–12072.

(48) Rodríguez-Pulido, A.; Kondrachuk, A. I.; Prusty, D. K.; Gao, J.; Loi, M. A.; Herrmann, A. Light-Triggered Sequence-Specific Cargo Release from DNA Block Copolymer–Lipid Vesicles. Angew. Chem. Int. Ed. 2013, 52, 1008–1012.

(49) Khmelinskaia, A.; Franquelim, H. G.; Petrov, E. P.; Schwille, P. Effect of Anchor Positioning on Binding and Diffusion of Elongated 3D DNA Nanostructures on Lipid Membranes. J. Phys. D Appl. Phys. 2016, 49, 194001.

(50) Khmelinskaia, A.; Mücksch, J.; Petrov, E. P.; Franquelim, H. G.; Schwille, P. Control of Membrane Binding and Diffusion of Cholesteryl-Modified DNA Origami Nanostructures by DNA Spacers. Langmuir 2018, 34, 14921–14931.

(51) Arnott, P. M.; Joshi, H.; Aksimentiev, A.; Howorka, S. Dynamic Interactions between Lipid-Tethered DNA and Phospholipid Membranes. Langmuir 2018, 34, 15084–15092.

(52) Morzy, D.; Rubio-Sánchez, R.; Joshi, H.; Aksimentiev, A.; Di Michele, L.; Keyser, U. F. Cations Regulate Membrane Attachment and Functionality of DNA Nanostructures. J. Am. Chem. Soc. 2021, 143, 7358–7367.

(53) Chidchob, P.; Daniel, O.; Mccarthy, D.; Luo, X.; Li, J.; Howorka, S.; Sleiman, H. F. Spatial Presentation of Cholesterol Units on a DNA Cube as a Determinant of Membrane Protein-Mimicking Functions. J. Am. Chem. Soc. 2019, 141, 1100–1108.

(54) Singh, J. K. D.; Darley, E.; Ridone, P.; Gaston, J. P.; Abbas, A.; Wickham, S. F. J.; Baker, M. A. B. Binding of DNA Origami to Lipids: Maximizing Yield and Switching via Strand Displacement. Nucleic Acids Res. 2021, 49, 10835–10850.

(55) Asandei, A.; Mereuta, L.; Bucataru, I. C.; Park, Y.; Luchian, T. A Single-Molecule Insight into the Ionic Strength-Dependent, Cationic Peptide Nucleic Acids-Oligonucleotides Interactions. Chem. Asian J. 2022, 17, e202200261.

(56) Yoder, B. L.; Bisson, R.; Beck, R. D. Steric Effects in the Chemisorption of Vibrationally Excited Methane on Ni(100). Science 2010, 329, 553–556.

(57) Martin, S. F.; Clements, J. H. Correlating Structure and Energetics in Protein-Ligand Interactions: Paradigms and Paradoxes. Annu. Rev. Biochem. 2013, 82, 267–293.

(58) Guo, Z.; Chen, B. Y. Explaining Small Molecule Binding Specificity with Volumetric Representations of Protein Binding Sites. J. Bioinform. Springer International Publishing 2022, 17–45.

(59) Gutlederer, E.; Gruhn, T.; Lipowsky, R. Polymorphism of Vesicles with Multi-Domain Patterns. Soft Matter 2009, 5, 3303–3311.

(60) Weng, J.; Xiang, X.; Ding, L.; Wong, A. L.-A.; Zeng, Q.; Sethi, G.; Wang, L.; Lee, S. C.; Goh, B. C. Extracellular Vesicles, the Cornerstone of next-Generation Cancer Diagnosis? Semin. Cancer Biol. 2021, 74, 105–120.

(61) Allelein, S.; Medina-Perez, P.; Lopes, A. L. H.; Rau, S.; Hause, G.; Kölsch, A.; Kuhlmeier, D. Potential and Challenges of Specifically Isolating Extracellular Vesicles from Heterogeneous Populations. Sci. Rep. 2021, 11, 11585.

(62) Steinkühler, J.; Sezgin, E.; Urbančič, I.; Eggeling, C.; Dimova, R. Mechanical Properties of Plasma Membrane Vesicles Correlate with Lipid Order, Viscosity and Cell Density. Commun. Biol. 2019, 2, 337.

(63) Heberle, F. A.; Feigenson, G. W. Phase Separation in Lipid Membranes. Cold Spring Harb. Perspect. Biol. 2011, 3, a004630.

(64) Santamaria, A.; Batchu, K. C.; Fragneto, G.; Laux, V.; Haertlein, M.; Darwish, T. A.; Russell, R. A.; Zaccai, N. R.; Guzmán, E.; Maestro, A. Investigation on the Relationship between Lipid Composition and Structure in Model Membranes Composed of Extracted Natural Phospholipids. J. Colloid Interface Sci. 2023, 637, 55–66.

(65) Willms, E.; Cabañas, C.; Mäger, I.; Wood, M. J. A.; Vader, P. Extracellular Vesicle Heterogeneity: Subpopulations, Isolation Techniques, and Diverse Functions in Cancer Progression. Front. Immunol. 2018, 9, 738.

(66) Ratajczak, M. Z.; Ratajczak, J. Extracellular Microvesicles/Exosomes: Discovery, Disbelief, Acceptance, and the Future? Leukemia 2020, 34, 3126–3135.

(67) Shimoda, A.; Ueda, K.; Nishiumi, S.; Murata-Kamiya, N.; Mukai, S. A.; Sawada, S. I.; Azuma, T.; Hatakeyama, M.; Akiyoshi, K. Exosomes as Nanocarriers for Systemic Delivery of the Helicobacter Pylori Virulence Factor CagA. Sci. Rep. 2016, 6, 18346.

(68) Koo, K. M.; Mainwaring, P. N.; Tomlins, S. A.; Trau, M. Merging New-Age Biomarkers and Nanodiagnostics for Precision Prostate Cancer Management. Nat. Rev. Urol. 2019, 16, 302– 317.

(69) Douglas, S. M.; Marblestone, A. H.; Teerapittayanon, S.; Vazquez, A.; Church, G. M.; Shih, W. M. Rapid Prototyping of 3D DNA-Origami Shapes with CaDNAno. Nucleic Acids Res. 2009, 37, 5001–5006.

(70) Ke, Y.; Douglas, S. M.; Liu, M.; Sharma, J.; Cheng, A.; Leung, A.; Liu, Y.; Shih, W. M.; Yan, H. Multilayer DNA Origami Packed on a Square Lattice. J. Am. Chem. Soc. 2009, 131, 15903–15908.

(71) Zhang, T.; Hartl, C.; Fischer, S.; Frank, K.; Nickels, P.; Heuer-Jungemann, A.; Nickel, B.; Liedl, T. 3D DNA Origami Crystals. Adv. Mater. 2018, 30, 1800273.

(72) Singh, J. K. D.; Luu, M. T.; Berengut, J. F.; Abbas, A.; Baker, M. A. B.; Wickham, S. F. J. Minimizing Cholesterol-Induced Aggregation of Membrane-Interacting DNA Origami Nanostructures. Membranes 2021, 11, 950.

(73) Lanphere, C.; Offenbartl-Stiegert, D.; Dorey, A.; Pugh, G.; Georgiou, E.; Xing, Y.; Burns, J. R.; Howorka, S. Design, Assembly, and Characterization of Membrane-Spanning DNA Nanopores. Nat. Protoc. 2021, 16, 86–130.

(74) Stengel, G.; Zahn, R.; Höök, F. DNA-Induced Programmable Fusion of Phospholipid Vesicles. J. Am. Chem. Soc. 2007, 129, 9584–9585.

(75) Paez-Perez, M.; Russell, I. A.; Cicuta, P.; Di Michele, L. Modulating Membrane Fusion through the Design of Fusogenic DNA Circuits and Bilayer Composition. Soft Matter 2022, 18, 7035–7044.

(76) Morzy, D.; Tekin, C.; Caroprese, V.; Rubio-Sánchez, R.; Michele, L. Di; Bastings, M. M. C. Interplay of the Mechanical and Structural Properties of DNA Nanostructures Determines Their Electrostatic Interactions with Lipid Membranes. Nanoscale 2023, 15, 2849–2859.

(77) Hatzakis, N. S.; Bhatia, V. K.; Larsen, J.; Madsen, K. L.; Bolinger, P. Y.; Kunding, A. H.; Castillo, J.; Gether, U.; Hedegård, P.; Stamou, D. How Curved Membranes Recruit Amphipathic Helices and Protein Anchoring Motifs. Nat. Chem. Biol. 2009, 5, 835–841.

(78) Van de Linde, S.; Löschberger, A.; Klein, T.; Heidbreder, M.; Wolter, S.; Heilemann, M.; Sauer, M. Direct Stochastic Optical Reconstruction Microscopy with Standard Fluorescent Probes. Nat. Protoc. 2011, 6, 991–1009.

(79) Batchelor, G. K. An Introduction to Fluid Dynamics; Cambridge University Press, 2000.

(80) Yokota, S.; Kuramochi, H.; Okubo, K.; Iwaya, A.; Tsuchiya, S.; Ichiki, T. Extracellular Vesicles Nanoarray Technology: Immobilization of Individual Extracellular Vesicles on Nanopatterned Polyethylene Glycol-Lipid Conjugate Brushes. PLoS One 2019, 14, e0224091.

(81) Kim, D. N.; Kilchherr, F.; Dietz, H.; Bathe, M. Quantitative Prediction of 3D Solution Shape and Flexibility of Nucleic Acid Nanostructures. Nucleic Acids Res. 2012, 40, 2862–2868.

(82) Wang, Y.; Wang, S.; Chen, A.; Wang, R.; Li, L.; Fang, X. Efficient Exosome Subpopulation Isolation and Proteomic Profiling Using a Sub-ExoProfile Chip towards Cancer Diagnosis and Treatment. Analyst 2022, 147, 4237–4248.

(83) Willms, E.; Johansson, H. J.; Mäger, I.; Lee, Y.; Blomberg, K. E. M.; Sadik, M.; Alaarg, A.; Smith, C. I. E.; Lehtiö, J.; El Andaloussi, S.; Wood, M. J. A.; Vader, P. Cells Release Subpopulations of Exosomes with Distinct Molecular and Biological Properties. Sci. Rep. 2016, 6, 22519.

(84) Beales, P. A.; Kyle Vanderlick, T. Partitioning of Membrane-Anchored DNA between Coexisting Lipid Phases. J. Phys. Chem. B 2009, 113, 13678–13686.

(85) Bryant, G.; Abeynayake, C.; Thomas, J. C. Improved Particle Size Distribution Measurements Using Multiangle Dynamic Light Scattering. 2. Refinements and Applications. Langmuir 1996, 12, 6224–6228.

(86) Schneider, C. A.; Rasband, W. S.; Eliceiri, K. W. NIH Image to ImageJ: 25 Years of Image Analysis. Nat. Methods 2012, 9, 671–675.

(87) Dempsey, G. T.; Vaughan, J. C.; Chen, K. H.; Bates, M.; Zhuang, X. Evaluation of Fluorophores for Optimal Performance in Localization-Based Super-Resolution Imaging. Nat. Methods 2011, 8, 1027–1036.

(88) Cnossen, J.; Cui, T. J.; Joo, C.; Smith, C. Drift Correction in Localization Microscopy Using Entropy Minimization. Opt. Express 2021, 29, 27961–27974.

