## Supplementary Information for "DNA Origami – Lipid Membrane Interactions Defined at Single-Molecular Resolution"

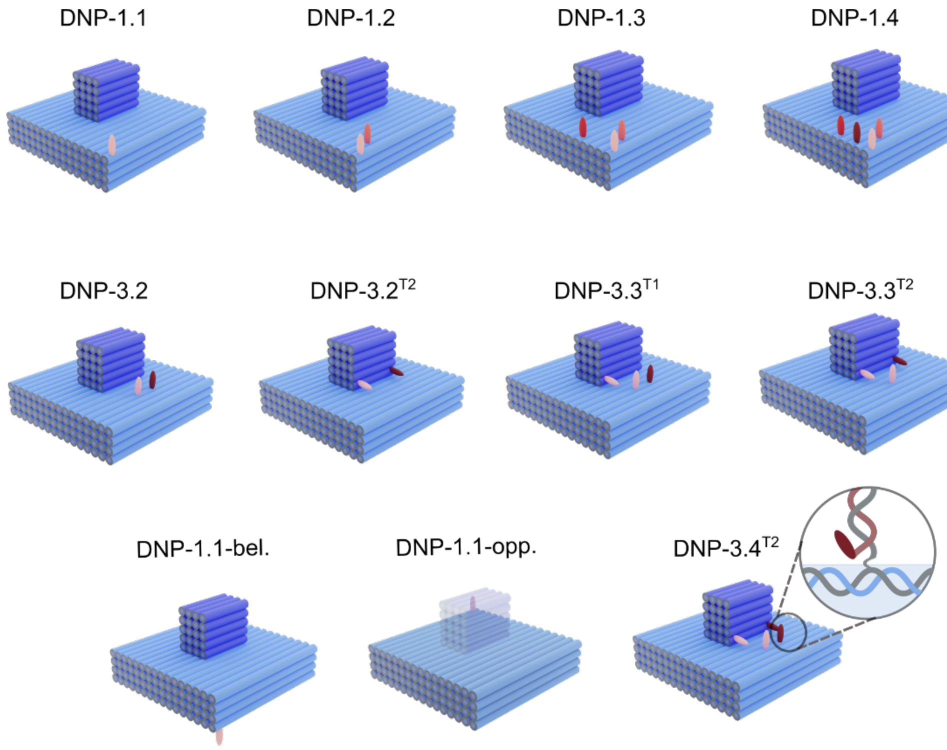


**Figure S1.** Schematic drawings of cholesterol-modified DNPs. DNP variants are named uses two digits. The first digit refers to the group and the second to the number of cholesterol moieties, as illustrated for group 1 and group 3 DNP-1 and DNP-3, respectively. In DNP-1 variants, cholesterol moieties are positioned at one corner of the DNP baseplate, while in DNP-3 versions cholesterols are positioned close to or on the tip of the DNP. For the latter, 'T' means that cholesterols are positioned at the DNP's tip as in DNP-3.2$^{T2}$ and DNP-3.3$^{T1}$ that carry two cholesterols and one cholesterol at the tip. In all structures, each cholesterol is color-coded to indicate its order of appearance within a group. DNP-1.1-bel carries a single cholesterol at the underside of the baseplate at a corner-edge position corresponding to DNP-1.1 while DNP-1.1-opp features a single cholesterol at the top side of the baseplate opposite of DNP-1.1.

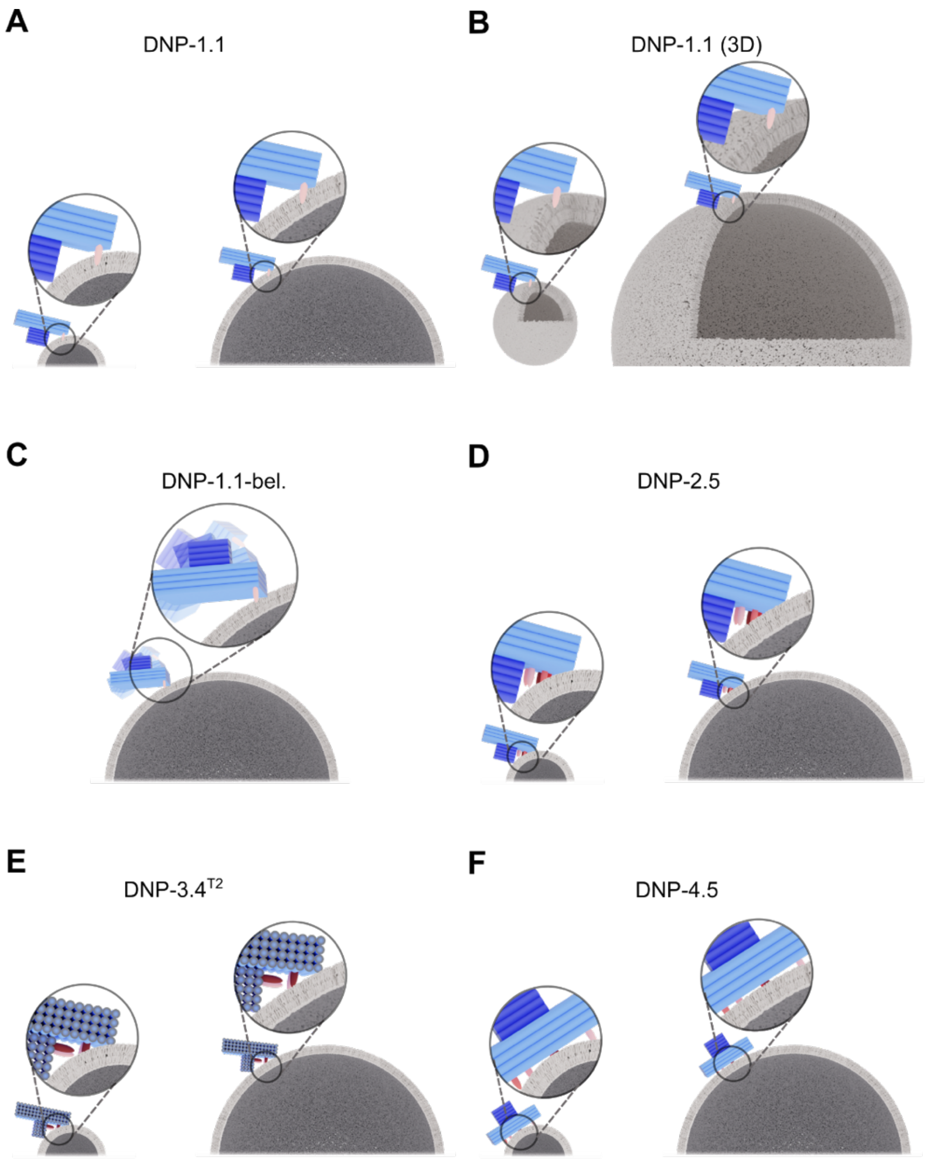


**Figure S2.** Schematic drawings on the geometrically defined interaction between various lipidated DNP and small and large unilamellar vesicles.

**Table S1.** Abbreviations, systematic chemical names, chemical structures, and phase transition temperatures of lipids used in this study. Phase transition temperatures are cited from Avanti Polar Lipids (https://avantilipids.com).

| **Lipid name** | **Lipid structure** | **Phase transition temperature (°C)** |
|---|---|---|
| DOPC<br><br>1,2-dioleoyl-*sn*-glycero-3-phosphocholine | 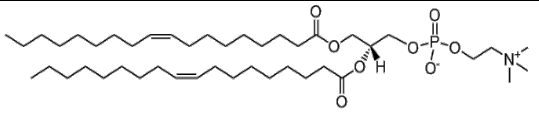 | -17 |
| DOPE<br><br>1,2-dioleoyl-*sn*-glycero-3-phosphoethanolamine | 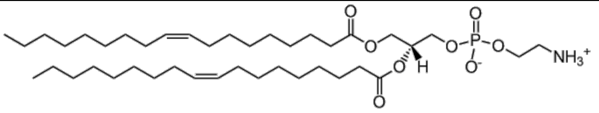 | -16 |
| POPC<br><br>1-palmitoyl-2-oleoyl-glycero-3-phosphocholine | 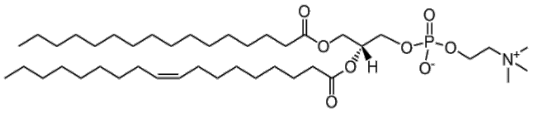 | -2 |
| Cy5-DOPE<br><br>1,2-dioleoyl-*sn*-glycero-3-phosphoethanolamine-N-(Cyanine 5) | 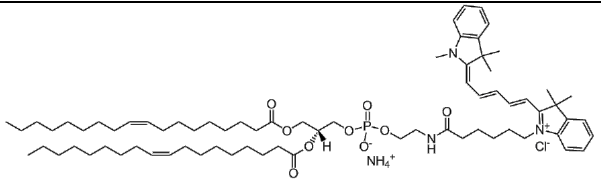 | N/A |
| Cholesterol | 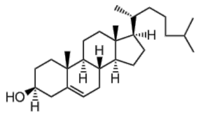 | N/A |
| Biotin-DOPE<br><br>1,2-dioleoyl-*sn*-glycero-3-phosphoethanolamine-N-(cap biotinyl) (sodium salt) | 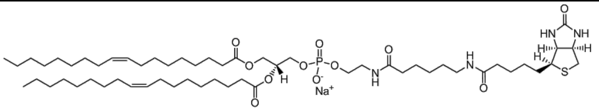 | N/A |

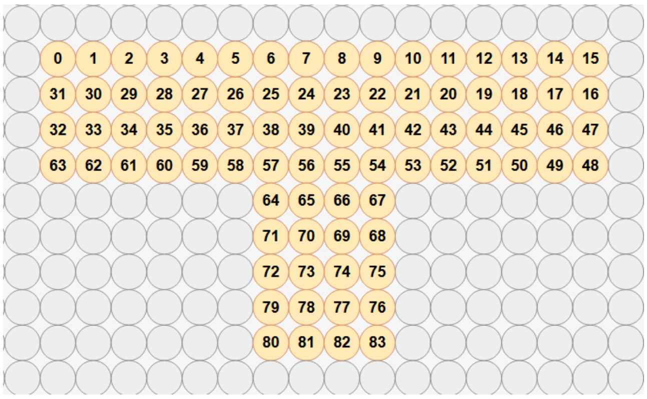


**Figure S3.** 2D duplex map of the DNA nanoprobe (DNP) indicating the position of the DNA helices. The T-shaped DNP is composed of a base plate of 16 parallel duplexes and four duplex layers high while the cuboid tip features 5 duplexes high and 4 duplexes wide.

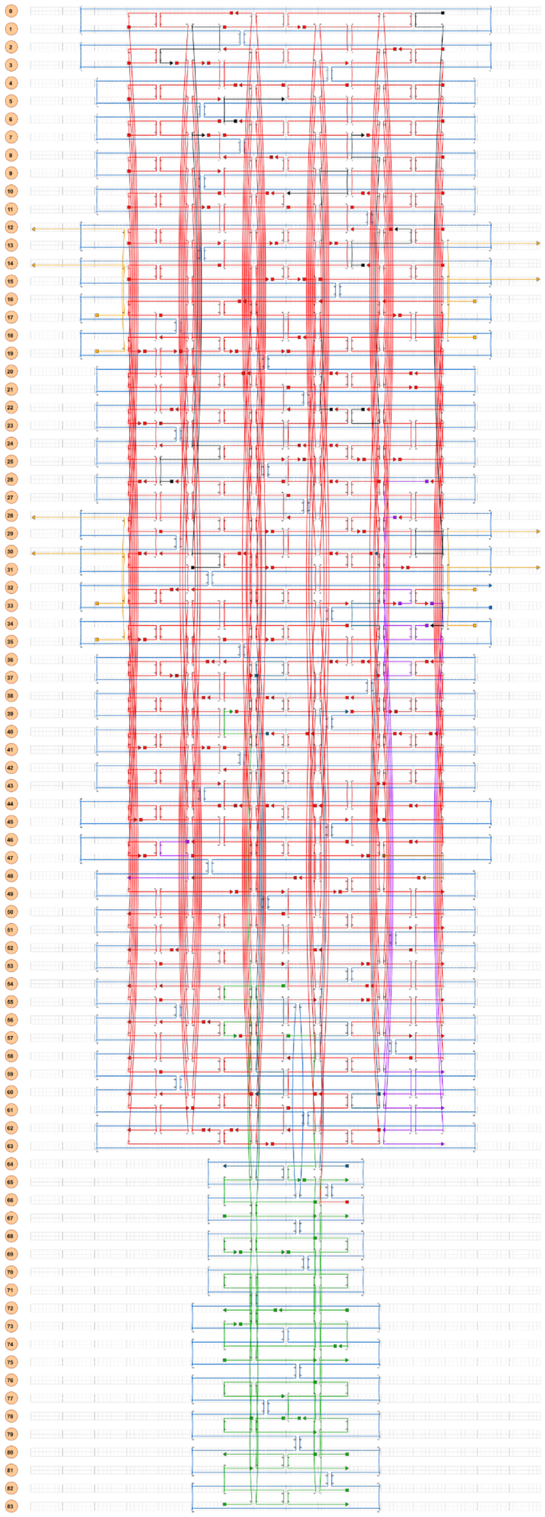

**Figure S4.** 2D design maps of DNP composed of a baseplate and a tip. The DNA strands are color-coded as followed. M13mp18 scaffold strand (blue), staple strands for baseplate (red); staple strands for tip (green); staple strands with handles for attachment of cholesterol-modified oligonucleotides for groups 1 and 2 DNP (pink), for group 3 DNP (dark blue), for group 4 (black); staple strands with handles for attachment of ATTO488-labelled oligos (gold).

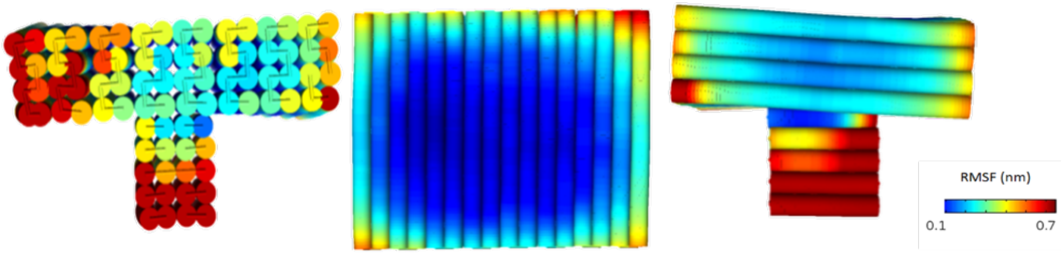


**Figure S5.** Result of simulation on expected thermal fluctuations of DNP performed in CanDo.[1] Front, bottom, and side view of the DNP as shown from left to right.

**Table S2.** Names and sequences of DNA staple strands used for assembling DNPs. Strand names annotated with * indicate strands that can be replaced by cholesterol handle homologous and cholesterol-modified oligonucleotides. Lowercase bases indicate the sequences of the handles used for binding cholesterol/fluorophore-modified oligonucleotides.

| **Position** | **Sequence (5'→3')** |
|---|---|
| Core*-1 | CAAAAGTTTTTTTTCAACTTTAATCATT |
| Core*-2 | TTGGAGAATAGCTACAGGTAGACGATAA |
| Core*-3 | AAATTAACCGTTGCTGATTGCAAGCGGTTACCAGACGAAAGATT |
| Core*-4 | GAAAGATTTTAAAACGAACT |
| Core*-5 | CATCAGTTAAATCTACTCCACTATGGACTCCAGC |
| Core*-6 | AGGAAAATCAAAAACAAGAGGTTAATAAGAACTGGC |
| Core*-7 | AAGCGTCAGAAAATCCATAGTAAGACATTCAA |
| Core*-8 | AATAAGTAATAAAAGATGATACCT |
| Core*-9 | TGACCAAAGACAAAAGGGCGCTCAATCGACAA |
| Core*-10 | AATCTAGCGATAGCTTAGAT |
| Core*-11 | GCACTTTGAATATCAGATGAATATACAG |
| Core*-12 | TCTGTAGCCGGAACTCATCTTTGACCCCTCCC |
| Core*-13 | AAACACAGTAAGCAAATATT |
| Core*-14 | TCTAGGGTACCGAGCTCGAAGGGTGCCTCATCGGAA |
| Core*-15 | GCCCCAGCAGGCTACATGGCGTGTCGAA |
| Core*-16 | GCGTAACGTTTTCTGTGCTAAACAACTTTCAA |
| Core*-17 | AAAAAGCCATAAGAATGGCGCTGGCAGAGCGGCAGT |
| Core-18 | CGCTTTTGAATTGAAACAAACATCAAGACTGATTGCAGAACGTA |
| Core-19 | TTTTGGAAGCTCAGTAATAAGTATTTTCCATT |
| Core-20 | CGAAACGCTAAAATATCTTTAGGAAACA |
| Core-21 | CTTGATCCTTATCATTCATCGAGATCTTCACA |
| Core-22 | ACCGCACTCCAAGAACACCAGCGCGGAA |
| Core-23 | AGGTGGTGAATTACGCATAACCGATATAAAAGACAGAATTGTAT |
| Core-24 | GCGGATAAGAAACGATGAGAGAATCGCTAATATCAGAGAGTAAA |
| Core-25 | AGGTGAGGCGGTCAGTAACCTCAAATGG |

| | |
|---|---|
| Core-26 | GCTCATACAGAGGAATTGCGTTGCGCTCATAAG |
| Core-27 | ATTAGAAAGCGCTAAATCCTAGGGAACCACTTTGAA |
| Core-28 | CTATTACGAGATGGGCAACCCGTCGGATTCTCACCGATAGTTGAGCCA |
| Core-29 | TACCAGTATTACGCCCTTTTACACAATAACGGATTCGCAAA |
| Core-30 | CAATCATTTTGCGGAACAAAAGAC |
| Core-31 | GCTTGACGGGGAAAAGCACTAGCCGCCAGACATCTT |
| Core-32 | GTCACCCTCAGCAGCGTTCGGTCGCCATCCCC |
| Core-33 | AGCAACAAAAAGCCTGTTCGTAATCATGGTCATGCCAAGC |
| Core-34 | GCACCTGCTTTAAACATGAGGAAGAGCC |
| Core-35 | CATTGAAACCACATTGAGGACCCTCAATCAATATCTGAT |
| Core-36 | CATTAAAAATACCGAAAATGGCTAGTTT |
| Core-37 | TTTCGAGCATGCGCCGGAGGCATTTAATAAGAGAATATAACAAT |
| Core-38 | ACCTCCCGTAAGAACGAGAACTGGAAGAAACGGGGT |
| Core-39 | GGATTATAATTTTTTCGCTTTCGACGAC |
| Core-40 | AGCAAAGAAGTGTTTTACTT |
| Core-41 | AGGAAGGGAACCAAATCAGGGC |
| Core-42 | CAACGCTCAACAGTAGGGCTTTAGGCACTAGACAG |
| Core-43 | GAGGACCCAGTAAGCAGGTCAGTTAAAATTAT |
| Core-44 | GCTGTCCCTTATACGGTACGCTT |
| Core-45 | AAGTTCACCGGATATTAGCGAATAAGGCTTGCCCTGACGAGAAATCAT |
| Core-46 | CATAGGCTGATACATAGGTCAGACGAT |
| Core-47 | GGTCACGAACGTGGATCATATGCGTTATACAAATTCTTAAAT |
| Core-48 | ATTCTTACCAAAACAACAG |
| Core-49 | CAAGGCGATTAAGTTGCGAGTAACGCATCGTA |
| Core-50 | CAAAAGGTTATCCTGTAGCAAACCAGGCAAGAT |
| Core-51 | CGGGATACCAAGATCCGCGACCTGCTCCTACGAGGCTGTGCCTT |
| Core-52 | CTAATGCAGGCTGACCAGCCACCAGAACCGCCCGC |
| Core-53 | CGGACACCGTACTCAGGAGGCAGAACCGCTGA |
| Core-54 | TTCAGGCTGCGCAACTTTTGAG |
| Core-55 | CTGAAATAAGTTAGAGGCTGGAATCGTCTGATTGTA |
| Core-56 | GCGTATGAGACGGTTGAGAAG |
| Core-57 | AATCAGATCATTACCGTAAG |
| Core-58 | GTAAAACGACGGCCAGTAGCTGTT |
| Core-59 | TACCCACCACCGCCCTCAGAGCCGCCACTTGAGGCAACGGACCA |
| Core-60 | GAGGCAGAACCAGTGTACAGCGTAACAA |
| Core-61 | AGTAAGTATGTTGAAAATACTGAGCTAAAAGG |
| Core-62 | CCACCTATTTTGCAATTTTATCCTGAATATCAGATGATAT |
| Core-63 | TAAAATCGGCCAACGCGCTCCT |
| Core-64 | TGCGAACAGCCCTGAGTTTCGTCACCAGTCTGC |
| Core-65 | ACCTTTTTTCAAAATCACGC |
| Core-66 | TCAAGTTAGAATCAAGTGTAACGT |
| Core-67 | GTACTGGTGTAGAAGAAAAA |
| Core-68 | CACGCCTCAGGAAAGCGCCATTC |
| Core-69 | GCAATTTTCATCGTAGGAATATAGAAGGACAT |

| | |
|---|---|
| Core-70 | CAAAAGGCAGATTTACCTACAAAGTATTATTAA |
| Core-71 | TCCTGTGTATACGAGCCCAATAGGTTGTTAAATCA |
| Core-72 | AGAGGCGGCAACAGGAACT |
| Core-73 | ATTACCTGCGCAGAGGTTCTGACCTGAAAGCGCTGCCCGCATTA |
| Core-74 | TAACAAGCATCACCTTGCTGATTAACACTATAGCTA |
| Core-75 | GGGAAACACCCTGAACAAAGGTGCCGGATTCCACAGATAGACCG |
| Core-76 | TTTTAGCGTCTTTCCAGAGCCCCAATCCGGCGCAAG |
| Core-77 | GCTTACCCCGATTAAAGC |
| Core-78 | GATTGTCAGTGCGTTCCAGT |
| Core-79 | GGGACGACGACAGTATCGGTTGGTGTCCAGCTGGATGTGCTG |
| Core-80 | GGCCTTGCTGGTTTCACCAGTTGGGCGCAATGTTTAGTTAATAT |
| Core-81 | AAATCAGAGCCCATGTACCAATCTCCAAAGGAACAACTAAAGGAAT |
| Core-82 | ATAACATTGAGGGAGGTTTGGGAAAACA |
| Core-83 | AACGCCAAGTTACAAAATCGAGCAAAAGAAGGGACAAGCCATTGTTT |
| Core-84 | TATCACCAGAAACCCTAAAGGGGTCAT |
| Core-85 | CCTTTACATTTTTGTTCCCAATAAAAACAATGTAG |
| Core-86 | GAGGGTGAAAGGACAGAAGGACATCATATTCCTG |
| Core-87 | GAACCACCACCCTTTAATTGTTGCAGGGA |
| Core-88 | ATTCATTGAGCCAGAAGGTAACTTAAACA |
| Core-89 | ATGGGCACCGCTTCTGTCA |
| Core-90 | ACCGTAACCACCGAGCACGTATAACGTGCCAGAATCTATTATAT |
| Core-91 | TTATTCATCAATATAATCCTACCTA |
| Core-92 | GCAGCCCTTCACAGGCTTTTTTTTGCCA |
| Core-93 | TGGGTCAACAATAGATTAAGACGCGATTAGTATTGC |
| Core-94 | GAGAAAACTGATGCAAGGGATTTTACCGAGTAAGTT |
| Core-95 | GAGTAGTTGGGAAGAAGAGATTTA |
| Core-96 | AGTAACAGTGCCGGAACCTAAAAAACCT |
| Core-97 | ATTAGTTGCAGCAGAAGATAAAAAGT |
| Core-98 | ATAGGGAGATACATAATTTACATTTTATTTTA |
| Core-99 | CACCCCAAAATTATTTATTTTCAGTTAGTCTT |
| Core-100 | GTAGCAATTATAATCAGCTTAGGTAACTATATGTAAATGCTTTT |
| Core-101 | CCACGCCTGCAACAGTGCCACGACGTT |
| Core-102 | ATCGGATCCGGTATTCACTTGCGGCGAGGAAATTAAG |
| Core-103 | CCTCTTATTCTGGGGGTTTTATGGATTAAGGTGGCACTTGAGAA |
| Core-104 | CATGATTTTGACGACATTCAATTAG |
| Core-105 | CGCGACCGTGTGTAGTTAATTTCATCTTCGCCTGTTCTGAATCA |
| Core-106 | TTATTTATCTAATTTGAGCC |
| Core-107 | ATCAATAATAATTTACTATCACCGAACGTCACACAG |
| Core-108 | GGAATACCAGCAACACTATCATAAGATAAATTTTTTGATGATACAACA |
| Core-109 | TTGCATGCACAAACTACAAACGGCTAATGGGATAGGT |
| Core-110 | AAACAGGCCGGATCACCGACCAAACAAAAGTC |
| Core-111 | TAATGCGCCACACAACGAAATTGTCTGGCCTTGCTTTCAT |
| Core-112 | AGTGAATAACAGTACATTACCGCCTTCTGGCCCCA |
| Core-113 | CAAACTCTGTAAAGTCAATAGTGAATTTAAACCTCCGGTGAGGCCAGAC |

| | |
|---|---|
| Core-114 | GGGTTAGAGATTGTTTAAAATCTAAACTAATACCTCTTCG |
| Core-115 | CTTTCGCAATAAGCCACCCTTTTAGTACGCGGGATCAAACGGGT |
| Core-116 | GCAAATATCATCGCCTCCCTCGTTCCAC |
| Core-117 | TCATTATAGGTTTAATGGGGTCGAGGTGCCGTAAGCCGGCGCTG |
| Core-118 | GTAGCCAGGGCGCGTACATTAGCAAACA |
| Core-119 | CTTTTGAGAAGATCGTCGCTATTAATTACAATTTCACATG |
| Core-120 | GCCGTATTACATTTAAATTTTCCCCCGATCAC |
| Core-121 | TAGTAAATTGGGCTTGAGATCCAGTCAGCCCAACGT |
| Core-122 | TGACACACCCGCTCCCTCAGAAATCGGACGAG |
| Core-123 | TGTTAAATAATATCCCATCCTCGGCTGTATAT |
| Core-124 | TAAGAAAATCAACAGTTAATTGAGAACATAAAAACAGGGAAGCG |
| Core-125 | GAATTTTACAAACAATTCGACAACGTTTGAGTGCAC |
| Core-126 | GATTGTTGACCATTACCTATGGTTAATGCCAG |
| Core-127 | CAAGAGCTCACATTCTTTGAGGCGAGGGTAGCAACGGCTTTT |
| Core-128 | TTGCGTAGGCACGTAAAACATCGC |
| Core-129 | TTAACGGAAGCATAAAGTGTTATTTTTGCGAA |
| Core-130 | TTTTTAATGGAAACCTTGCTTATC |
| Core-131 | CGCACTCCAGCCAGCTTTCCGGATTTT |
| Core-132 | CCTGTCGCCGACAATGACAACAAC |
| Core-133 | AGCCGCGCTAACGGAAGCCTTAAATCAAGTAAC |
| Core-134 | AGAACAGCTAATGCAGAACGCTGACCTAGCTT |
| Core-135 | TGATCCAGGCGGCTACGAAGAAAACGAA |
| Core-136 | GAACTCCAAAAGGAATATGTTACTAATTTACCCTTG |
| Core-137 | TGAAATCTAAAGAGAATAGAAAAAAAAG |
| Core-138 | GGCTATCGGTTTATCAGCTTACGTTGAAGTAACACTCATAGTTA |
| Core-139 | TTTAATCAATAGCCTATTTCCGTATAAAACTAAAACACGAGGCG |
| Core-140 | TCCTCATATATTTATAAATAAGGCGTTAATGTTTAGTCGAGAA |
| Core-141 | CCCTACCAGAGCCAAATCAAACCAGGCG |
| Core-142 | GCTGCTTTCCAGTTTAAATTTAACGCCAT |
| Core-143 | CATCGCCCTCTTAAACAGCTTGATCGTGGGAACAAGGGGG |
| Core-144 | AAGTCCTGTTAGAGCCCCAGTAGCATGG |
| Core-145 | TTCGATTAATGGTTTGACCAGTATTAGA |
| Core-146 | TTTTAAAATCGTATTACATTTGAGGATTTAGA |
| Core-147 | TGGTACCCTCAGTTCATCAAGACAAGAACCGGA |
| Core-148 | GAAACACCAGTCACACGACCCGTG |
| Core-149 | ACATGTAATTAATTGAGCGTTTTCATCGGCATAGCC |
| Core-150 | TAAAGTCAGAACCGCCTCCGAAATCGGCAAAAGTTT |
| Core-151 | GACGACGAAGTACCGACGTAATCAAATC |
| Core-152 | ATAACATGATTAAGACTCCTAAGGAAACGAGG |
| Core-153 | GCCAAGTATTAGACTAACTG |
| Core-154 | ATGATTGTAAACGACTGGATAGCGTCCAACGGAGATGCAGGAGT |
| Tip*-1 | ACGGTAATAAAGTACAATACTGCGAGACGGGG |
| Tip*-2 | ATTAAACGAGAAATAAATATATCAGAAATAGCATGTCAATCATA |
| Tip-3 | AGTAATTTTGAGAAAGGCTATCCTGAGAG |

| | |
|---|---|
| Tip-4 | TTAAGCAAGAGCATAACCAAAAACCAAAAGGGCTG |
| Tip-5 | CAGTCAAATGATAAATGAGAGGGTAAAAGAGA |
| Tip-6 | TGGCATCATATAATGCTTAAATATGCAAATAC |
| Tip-7 | AAAGTACGTTCCATATGACCGGAA |
| Tip-8 | TAAAGCCCAATGGTCAATACAAGG |
| Tip-9 | ATTATAATTCGAGCGAACCAATTCAACCGTTCTAGCTCACCATCAA |
| Tip-10 | ACAGGACCGAGAAGCCGATAAAAACCTCATAT |
| Tip-11 | CGCGAAACCGTAAAACAGCC |
| Tip-12 | CAGGTGCAAAGCGGATGGCTTAGAGCTAGAAC |
| Tip-13 | ACTTCTATTGTGTAGGCTTTAATTGCTCCTTTCAGG |
| Tip-14 | TTGCGGATTGCATCAAAGAG |
| Tip-15 | TATGATAACAGTTGATTCCTCA |
| Tip-16 | GAAGACCATAAATCAACCCGAAAG |
| Tip-17 | CCAAAGCAAATATCGCGTTTGAGAGTACTAAAGATTATTA |
| Tip-18 | TGACTTTTGCGGTGTTTAGCAATAGTAGTAGCATTA |
| Tip-19 | CTGACTATCTTTTTCAGTTCAGAAGGATTAGCAAA |
| Tip-20 | ACTAAAGATATAGTCAGAACATTG |
| Tip-21 | TTTTTTAATTGCTGAAATTCTACTTATATTTTCATTTGGGTTAGATAC |
| Tip-22 | CCTGTACTATTTCGCAAATTCTGCGAACGAGT |
| Tip-23 | GAAGAAATCAGGGGTTGATATCATTGAACAGC |
| Core-Fluoro.-1 | CAATAGATAATAAATCCTTT tttagctcactcagtctca |
| Core-Fluoro.-2 | TGAGTGTTGTTCGAGCTAAA tttagctcactcagtctca |
| Core-Fluoro.-3 | GTGAACCATCACGAAAGCGA tttagctcactcagtctca |
| Core-Fluoro.-4 | AATTGAGTTAAGTAACGTCA tttagctcactcagtctca |
| Core-Fluoro.-5 | CCGCCAGCATTGCAATGAAA tttagctcactcagtctca |
| Core-Fluoro.-6 | CTATCTTACCGACCAGTTAC tttagctcactcagtctca |
| Core-Fluoro.-7 | TCATAATCAAAATTGCCTTT tttagctcactcagtctca |
| Core-Fluoro.-8 | AGCAGCAAATGAGGATTATA tttagctcactcagtctca |
| DNP-1.1 / 2.5 | CAAAAGTTTTTTTTCAACTTTAATCATT tttcctacgagtcagtcc |
| DNP-1.2 / 2.2 | AGGAAAATCAAAAACAAGAGGTTAATAAGAACTGGC tttcctacgagtcagtcc |
| DNP-1.3 / 2.3 | TTGGAGAATAGCTACAGGTAGACGATAA tttcctacgagtcagtcc |
| DNP-1.4 / 2.4 | GAAAGATTTTAAAACGAACT tttcctacgagtcagtcc |
| DNP-2.1 | AAATTAACCGTTGCTGATTGCAAGCGGTTACCAGACGAAAGATT tttcctacgagtcagtcc |
| DNP-3.2$^{T0}$ | AAGCGTCAGAAAATCCATAGTAAGACATTCAA tttcctacgagtcagtcc |
| DNP-3.2$^{T0}$ | catcttcacactactttt CATCAGTTAAATCTACTCCACTATGGACTCCAGC |
| DNP-3.3$^{T1}$ | ATTAAACGAGAAATAAATATATCAGAAATAGCATGTCAATCATA tttcctacgagtcagtcc |
| DNP-3.4$^{T2}$ | catcttcacactactttt ACGGTAATAAAGTACAATACTGCGAGACGGGG |
| DNP-4.1 | AATAAGTAATAAAAGATGATACCT tttcctacgagtcagtcc |
| DNP-4.2 | TGACCAAAGACAAAAGGGCGCTCAATCGACAA tttcctacgagtcagtcc |

| | |
|---|---|
| DNP-4.3 | TTTCGAGCATGCGCCGGAGGCATTTAATAAGAGAATATAACAAT tttcctacgagtcagtcc |
| DNP-4.4 | AATCTAGCGATAGCTTAGAT tttcctacgagtcagtcc |
| DNP-4.5 | GCACTTTGAATATCAGATGAATATACAG tttcctacgagtcagtcc |
| DNP-1.1-bel. | GCGTAACGTTTTCTGTGCTAAACAACTTTCAA tttcctacgagtcagtcc |
| DNP-1.1-opp. | catcttcacactactttt AAAAAGCCATAAGAATGGCGCTGGCAGAGCGGCAGT |

**Table S3.** List of oligonucleotides carrying fluorophore and cholesterol-modified oligonucleotides.

| **Name** | **Sequence** |
|---|---|
| Oligo-ATTO488-3' | TGAGACTGAGTGAGCT - ATTO488 |
| Oligo-Cholesterol- 5' | Cholesterol - TEG - AGTAGTGTGAAGATG |
| Oligo-Cholesterol- 3' | GGACTGACTCGTAGG - TEG - cholesterol |

**Table S4.** Annealing protocol for the DNP depicting the different steps and time for each step along with the initial annealing temperature ($T_i$) and the final annealing temperature ($T_f$).

| **Step** | **$T_i$ (C°)** | **$T_f$ (C°)** | **Time (min)** |
|---|---|---|---|
| 1 | 95 | 95 | 1 |
| 2 | 85 | 65.1 | 16 |
| 3 | 65 | 59.1 | 300 |
| 4 | 59 | 54.1 | 525 |
| 5 | 54 | 45.1 | 450 |
| 6 | 45 | 41.1 | 420 |
| 7 | 41 | 36.1 | 250 |
| 8 | 36 | 4 | 26 |
| | | **TOTAL TIME:** | 1988 |

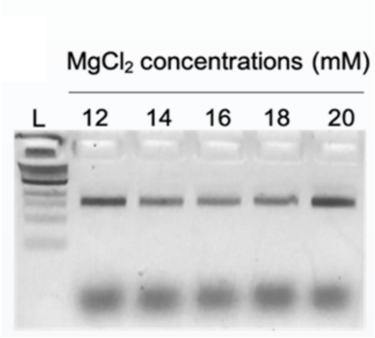


**Figure S6.** DNP assembly confirmed by 1.5% agarose gel electrophoresis. DNP was assembled in TAE supplemented with $MgCl_2$ ranging in concentration from 12 - 20 mM. L, 1 kb DNA ladder.

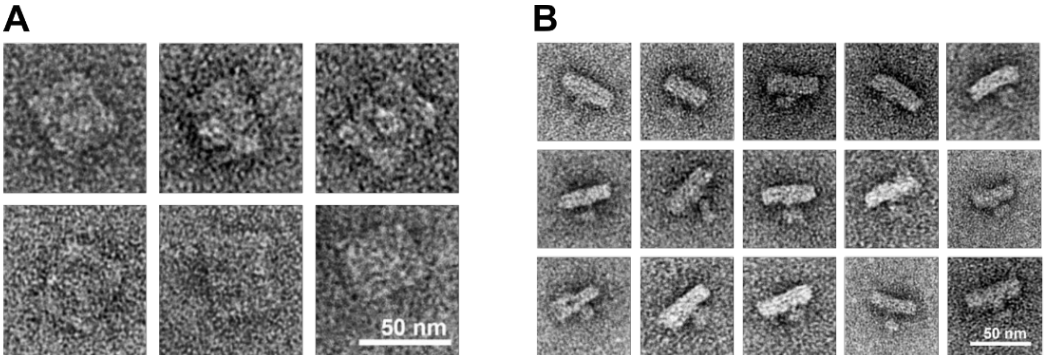


**Figure S7.** TEM images of negatively stained DNP. (A) Bottom and top view, (B) side views of DNPs.

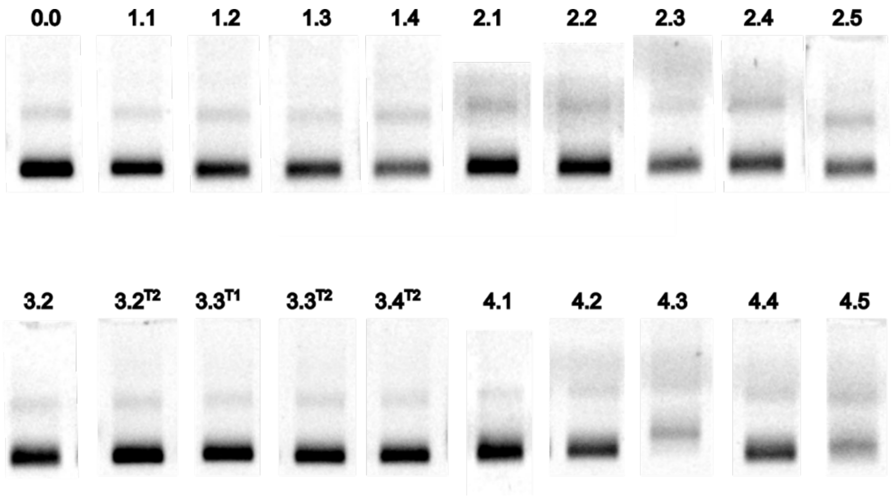


**Figure S8.** 2% Agarose gel electrophoretic analysis of purified lipidated DNPs. The two digits identifier refers to the naming of DNP, as defined in Figure S6. Gel lanes for DNP-4.3 and DNP-4.5 indicate a lower yield of monomeric DNP due to hydrophobically induced dimerization and aggregation.

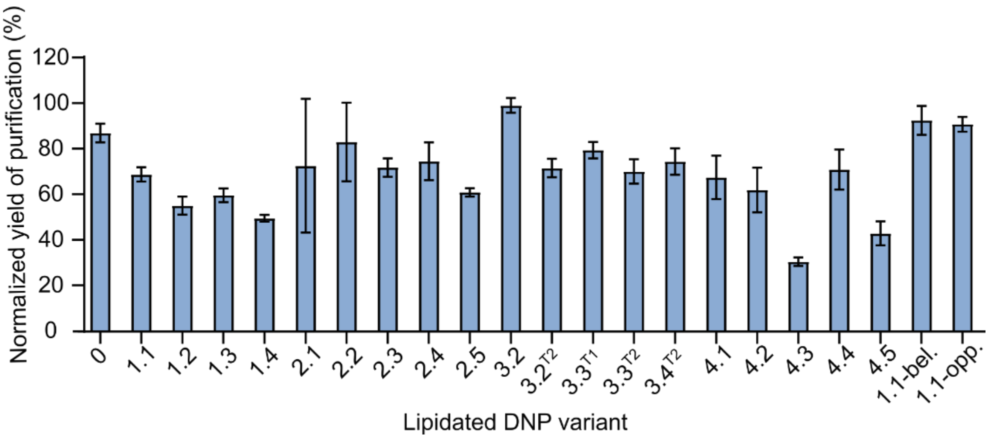


**Figure S9.** Yields of lipidated DNPs after purification via spin-column filtration. Data were obtained from gel band intensities and normalizing so 100% corresponds to a DNP condition folded without cholesterols. For comparison between gels, gel band intensities were normalized with four only-scaffold DNA controls included in each gel as an internal calibration standard. The data represent the averages and standard error of the mean for at least 3 replicas of each experiment.

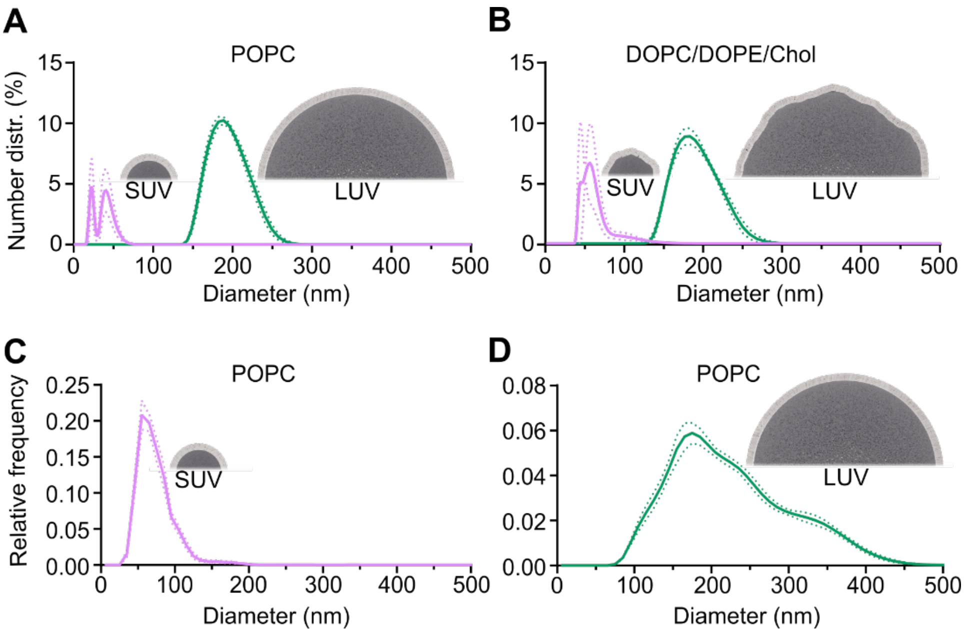


**Figure S10.** Size distribution analysis of small (SUVs) and large (LUVs) unilamellar vesicles determined by dynamic light scattering (DLS) and nanoparticle tracking analysis (NTA). (A,B) DLS analysis of (A) SUVs and LUVs with POPC bilayers and (B) with DOPC/DOPE/Chol bilayers with a molar ratio 0.4:0.2:0.4. The weighted means of the diameter are (A) 34 ± 4 nm and 192 ± 2 nm, and (B) 58 ± 4 nm and 187 ± 2 nm. (C,D) NTA analysis of relative frequency against diameter of (C) SUVs and (D) LUVs with POPC bilayers. The weighted means of the diameter are 75 ± 2 nm and 225 ± 12 nm. Solid and dotted lines represent average and standard error of the mean.

**Table S5.** Quantitative results on DNP binding to vesicles obtained from gel shift analysis. The data include the original gel band intensities for each experiment, calculated relative average binding, and standard error for binding to POPC and DOPC/DOPE/Chol SUVs and LUVs.

| **POPC SUVs** | Experiment | | | | | | | | | |
|---|---|---|---|---|---|---|---|---|---|---|
| **DNP-** | **1** | **2** | **3** | **4** | **5** | **6** | **7** | **Average, unbound DNP (%)** | **Error (%)** | **Average, bound DNP (%)** |
| **0** | 0.95 | 1.05 | 0.95 | 1.02 | 1.00 | | | 100 | 2 | 0 |
| **1.1** | 0.85 | 1.06 | 0.94 | 0.86 | 0.80 | 1.14 | 0.96 | 93 | 4 | 7 |
| **1.1-Unfold Tip** | 0.91 | 0.20 | 0.94 | 1.02 | | | | 97 | 3 | 3 |
| **DNP-1.1-bel.** | 0.93 | 1.00 | 0.85 | | | | | 93 | 4 | 7 |
| **DNP-1.1-opp.** | 0.92 | 0.96 | | | | | | 94 | 2 | 6 |
| **1.2** | 0.25 | 0.20 | 0.28 | 0.28 | | | | 25 | 2 | 75 |
| **1.3** | 0.18 | 0.18 | 0.26 | 0.25 | | | | 22 | 2 | 78 |
| **1.4** | 0.50 | 0.28 | 0.21 | 0.39 | 0.40 | | | 36 | 5 | 64 |
| **0-** | 0.98 | 0.99 | 1.03 | 1.05 | 1.04 | | | 102 | 1 | -2 |
| **2.1** | 0.98 | 0.41 | 0.48 | 0.95 | 1.01 | | | 76 | 13 | 24 |
| **2.2** | 0.15 | 0.50 | 0.43 | | | | | 36 | 11 | 64 |
| **2.3** | 0.57 | 0.38 | 0.49 | 0.49 | 0.39 | | | 46 | 4 | 54 |
| **2.4** | 0.32 | 0.39 | 0.46 | 0.50 | | | | 42 | 4 | 58 |

| | | | | | | | | | | |
|---|---|---|---|---|---|---|---|---|---|---|
| **2.5** | 0.17 | 0.23 | 0.31 | 0.36 | | | | 27 | 4 | 73 |
| **3.2** | 0.63 | 0.57 | 0.82 | 0.86 | 0.87 | 0.83 | | 76 | 5 | 24 |
| **3.2$^{T2}$** | 0.42 | 0.78 | 0.45 | 0.80 | 0.76 | 0.53 | 0.42 | 62 | 7 | 38 |
| **3.3$^{T1}$** | 0.47 | 0.57 | 0.76 | 0.70 | | | | 62 | 7 | 38 |
| **3.3$^{T2}$** | 0.42 | 0.74 | 0.66 | 0.64 | | | | 62 | 7 | 39 |
| **3.4$^{T2}$** | 0.43 | 0.30 | 0.57 | 0.56 | | | | 46 | 7 | 54 |
| **4.1** | 1.05 | 1.09 | | 1.00 | 1.03 | | | 104 | 2 | -4 |
| **4.2** | 1.05 | 0.99 | 1.00 | 1.05 | | | | 102 | 2 | -2 |
| **4.3** | 0.35 | 0.61 | 0.72 | 0.74 | 0.84 | | | 65 | 8 | 35 |
| **4.4** | 0.58 | 0.49 | 0.57 | 0.80 | 0.66 | | | 62 | 5 | 38 |
| **4.5** | 0.08 | 0.29 | 0.32 | 0.51 | 0.39 | | | 32 | 7 | 68 |
| **POPC LUVs** | | | | | | | | | | |
| **DNP-** | **1** | **2** | **3** | **4** | **5** | **6** | **7** | **Average, unbound DNP (%)** | **Error (%)** | **Average, bound DNP (%)** |
| **0** | 0.92 | 1.13 | 0.94 | 0.98 | 0.94 | | | 98 | 4 | 2 |
| **1.1** | 0.72 | 0.33 | 0.56 | 0.49 | 0.47 | 0.33 | 0.32 | 45 | 5 | 55 |
| **1.1-Unfold Tip** | 0.61 | 0.73 | 0.65 | 0.70 | | | | 67 | 3 | 33 |
| **DNP-1.1-bel.** | 0.91 | 0.90 | 0.83 | | | | | 88 | 2 | 12 |
| **DNP-1.1-opp.** | 0.55 | 0.42 | | | | | | 48 | 6 | 52 |
| **1.2** | 0.16 | 0.20 | 0.24 | 0.24 | | | | 21 | 2 | 79 |
| **1.3** | 0.25 | 0.16 | 0.20 | 0.19 | | | | 20 | 2 | 80 |
| **1.4** | 0.22 | 0.27 | 0.27 | 0.29 | 0.31 | | | 27 | 2 | 73 |
| **0-** | 0.94 | 1.02 | 0.99 | 1.09 | 1.06 | | | 102 | 3 | -2 |
| **2.1** | 1.05 | 0.94 | 0.49 | 0.54 | 0.86 | 0.98 | | 81 | 10 | 19 |
| **2.2** | 0.72 | 0.19 | 0.40 | 0.45 | | | | 44 | 11 | 56 |
| **2.3** | 0.71 | 0.42 | 0.36 | 0.38 | 0.44 | | | 46 | 6 | 54 |
| **2.4** | 0.47 | 0.39 | 0.38 | 0.40 | | | | 41 | 2 | 59 |
| **2.5** | 0.26 | 0.22 | 0.20 | 0.23 | | | | 23 | 1 | 77 |
| **3.2** | 1.03 | 0.63 | 0.86 | 0.88 | 0.91 | 0.93 | | 87 | 5 | 13 |
| **3.2$^{T2}$** | 0.90 | 0.73 | 0.69 | 0.73 | 0.77 | | | 76 | 4 | 24 |
| **3.3$^{T1}$** | 0.67 | 0.55 | 0.57 | 0.59 | 0.88 | | | 65 | 6 | 35 |
| **3.3$^{T2}$** | 0.69 | 0.59 | 0.57 | 0.63 | | | | 62 | 3 | 38 |
| **3.4$^{T2}$** | 0.54 | 0.55 | 0.48 | 0.54 | | | | 52 | 2 | 48 |
| **4.1** | 0.87 | 0.99 | 1.01 | 1.08 | | | | 99 | 4 | 1 |
| **4.2** | 0.88 | 0.78 | 0.77 | 0.90 | | | | 83 | 3 | 17 |
| **4.3** | 0.21 | 0.47 | 0.37 | 0.48 | 0.71 | | | 45 | 8 | 55 |
| **4.4** | 0.09 | 0.39 | 0.26 | 0.49 | 0.53 | | | 35 | 8 | 65 |
| **4.5** | 0.09 | 0.25 | 0.19 | 0.27 | 0.38 | | | 23 | 5 | 77 |
| **DOPC/DOPE/Chol SUVs** | | | | | | | | | | |
| **DNP-** | **1** | **2** | **3** | **4** | **5** | | | **Average, unbound DNP (%)** | **Error (%)** | **Average, bound DNP (%)** |
| **0** | 1.00 | 0.95 | 0.99 | | | | | 98 | 1 | 2 |
| **1.1** | 0.68 | 0.47 | 0.53 | 0.61 | 0.64 | | | 58 | 4 | 42 |
| **1.2** | 0.13 | 0.18 | 0.21 | | | | | 17 | 2 | 83 |
| **1.3** | 0.10 | 0.16 | 0.15 | | | | | 14 | 2 | 86 |

| | | | | | | | | | | |
|---|---|---|---|---|---|---|---|---|---|---|
| **1.4** | 0.06 | 0.18 | 0.15 | | | | | 13 | 4 | 87 |
| **0-** | 1.00 | 1.04 | 1.02 | 0.72 | | | | 94 | 8 | 6 |
| **2.1** | 0.90 | 0.86 | 0.94 | 0.82 | | | | 88 | 3 | 12 |
| **2.2** | 0.28 | 0.35 | 0.38 | | | | | 34 | 3 | 66 |
| **2.3** | 0.26 | 0.32 | 0.34 | | | | | 31 | 2 | 69 |
| **2.4** | 0.13 | 0.19 | 0.21 | | | | | 17 | 2 | 83 |
| **2.5** | 0.05 | 0.11 | 0.15 | | | | | 10 | 3 | 90 |
| **3.2** | 0.72 | 0.79 | 0.93 | | | | | 81 | 6 | 19 |
| **3.2**$^{T2}$ | 0.56 | 0.78 | 0.81 | 0.62 | 0.54 | | | 66 | 7 | 34 |
| **3.3**$^{T1}$ | 0.39 | 0.54 | 0.59 | | | | | 50 | 6 | 50 |
| **3.3**$^{T2}$ | 0.34 | 0.59 | 0.60 | | | | | 51 | 9 | 49 |
| **3.4**$^{T2}$ | 0.21 | 0.55 | 0.58 | | | | | 44 | 12 | 56 |
| **4.1** | 0.96 | 1.01 | 0.91 | | | | | 96 | 3 | 4 |
| **4.2** | 0.69 | 0.74 | 0.78 | | | | | 73 | 3 | 27 |
| **4.3** | 0.12 | 0.19 | 0.23 | | | | | 18 | 3 | 82 |
| **4.4** | 0.14 | 0.16 | 0.15 | | | | | 15 | 1 | 85 |
| **4.5** | 0.06 | 0.24 | 0.22 | | | | | 17 | 6 | 83 |
| **DOPC/DOPE/Chol LUVs** | | | | | | | | | | |
| **DNP-** | **1** | **2** | **3** | **4** | **5** | | | **Average, unbound DNP (%)** | **Error (%)** | **Average, bound DNP (%)** |
| **0** | 0.94 | 1.08 | 0.93 | | | | | 98 | 5 | 2 |
| **1.1** | 0.41 | 0.38 | 0.37 | 0.29 | 0.35 | | | 36 | 2 | 64 |
| **1.2** | 0.14 | 0.13 | 0.16 | | | | | 14 | 1 | 86 |
| **1.3** | 0.08 | 0.13 | 0.11 | | | | | 10 | 1 | 90 |
| **1.4** | 0.09 | 0.12 | 0.11 | | | | | 11 | 1 | 89 |
| **0-** | 1.02 | 1.19 | 1.00 | 0.96 | | | | 104 | 5 | -4 |
| **2.1** | 0.75 | 0.89 | 0.88 | 0.85 | | | | 84 | 3 | 16 |
| **2.2** | 0.25 | 0.33 | 0.32 | | | | | 30 | 2 | 70 |
| **2.3** | 0.28 | 0.30 | 0.28 | | | | | 29 | 1 | 71 |
| **2.4** | 0.15 | 0.17 | 0.21 | | | | | 18 | 2 | 82 |
| **2.5** | 0.05 | 0.10 | 0.09 | | | | | 8 | 1 | 92 |
| **3.2** | 0.75 | 0.83 | 0.86 | | | | | 81 | 3 | 19 |
| **3.2**$^{T2}$ | 0.68 | 0.83 | 0.75 | 0.56 | 0.55 | | | 67 | 7 | 33 |
| **3.3**$^{T1}$ | 0.44 | 0.56 | 0.50 | | | | | 50 | 4 | 50 |
| **3.3**$^{T2}$ | 0.52 | 0.65 | 0.58 | | | | | 58 | 4 | 42 |
| **3.4**$^{T2}$ | 0.31 | 0.64 | 0.55 | | | | | 50 | 10 | 50 |
| **4.1** | 1.00 | 0.91 | 0.89 | | | | | 93 | 3 | 7 |
| **4.2** | 0.70 | 0.80 | 0.69 | | | | | 73 | 4 | 27 |
| **4.3** | 0.12 | 0.09 | 0.09 | | | | | 10 | 1 | 90 |
| **4.4** | 0.11 | 0.09 | 0.11 | | | | | 10 | 1 | 90 |
| **4.5** | 0.05 | 0.08 | 0.13 | | | | | 9 | 2 | 91 |

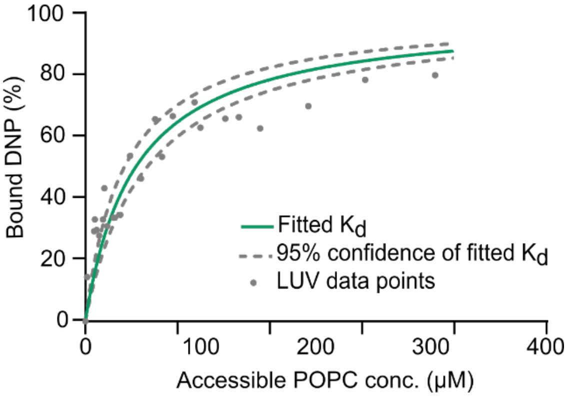


**Figure S11.** Plot on the binding of lipidated DNP-1.1 to POPC LUVs as a function of POPC concentration. The POPC concentration was obtained from the DLS data by assuming a packing density for POPC bilayers of 0.627 nm$^2$ per lipid molecule. The lipid area available for DNA nanostructure binding is always in excess thereby fulfilling the requirement for a Langmuir isotherm. The data from two different experiments were fitted to a Langmuir isotherm $Fraction\ bound = \frac{[POPC]}{[POPC]+Kd}$ to yield a $K_d$ of 5.4 ± 1.2 x 10$^{-5}$ M.

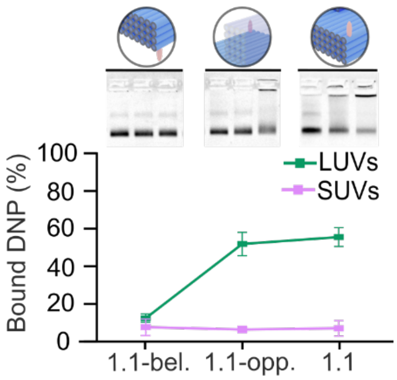


**Figure S12.** Binding of lipidated DNPs to POPC SUVs and LUVs depends on the cholesterol position and vesicle size. Agarose gel electrophoresis and quantitative analysis on the interaction for DNP-1.1-bel, DNP-1.1-opp, and DNP-1.1.

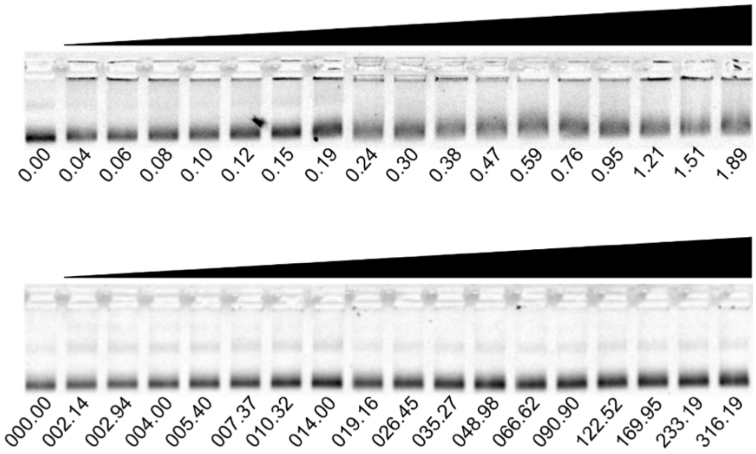


**Figure S13.** DNP-1.1 does not bind to POPC vesicles as shown by minimal gel band upshift in 2% agarose gel electrophoresis. DNP-1.1 was incubated with LUVs (top row) and SUVs (bottom row) for 15 h. The numbers below the gels correspond to the vesicle concentration values in nM.

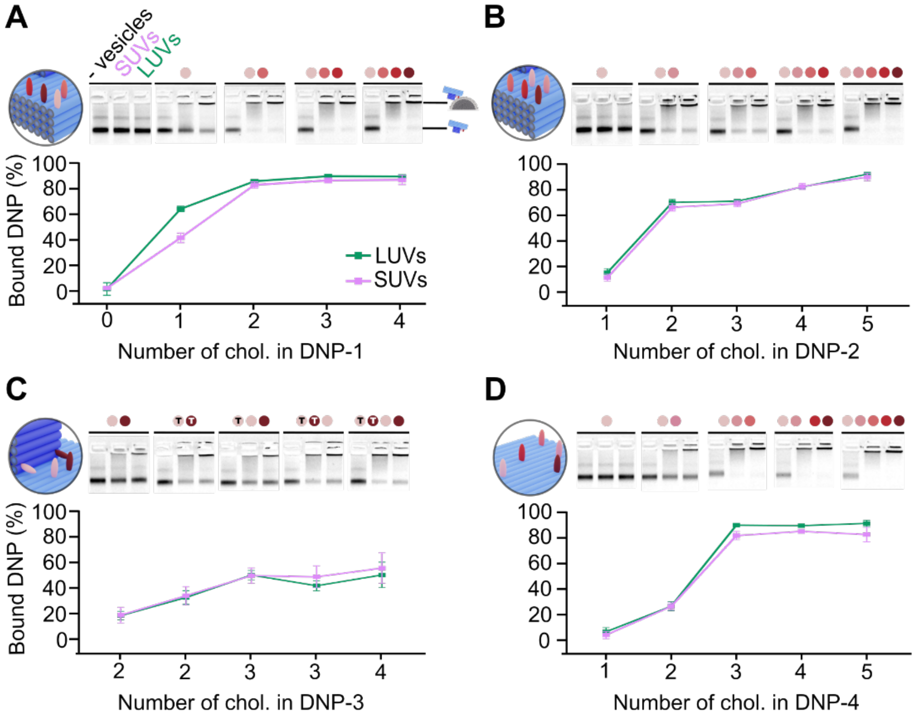


**Figure S14.** Binding of lipidated DNPs to DOPC/DOPE/Chol vesicles against cholesterol number and position, and vesicle size**.** Agarose gel electrophoresis and quantitative analysis on vesicle binding for DNPs of (A) DNP-1, (B) DNP-2, (C) DNP-3, and (D) DNP-4. The agarose gels show each three lanes for free DNP, DNP after SUV incubation, and DNP after LUV incubation, as linked together with a horizontal line. The number and position of cholesterols in DNPs are indicated by red-colored dots at the top of each three lanes. The color-coded cholesterols in DNP are schematically illustrated to the left of each gel in the panel. In C, the "T" on colored dots represents the location of cholesterol at the DNP's tip. The data points represent averages and the standard error from at least three independent experiments.

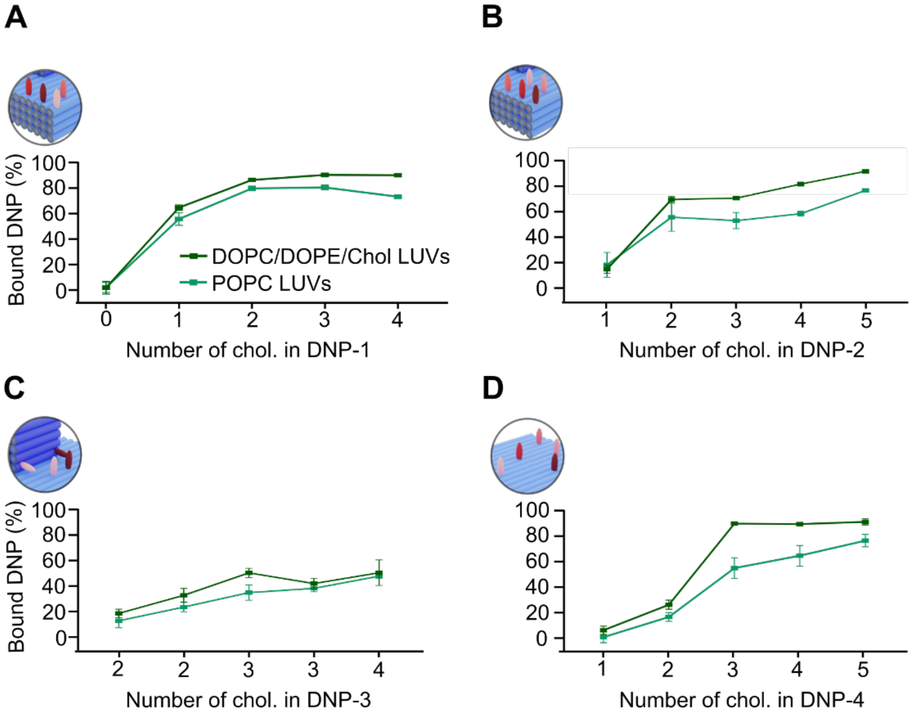


**Figure S15.** Binding of lipidated DNPs to DOPC/DOPE/Chol LUVs and POPC LUVs against cholesterol number and position, and vesicle size. Quantitative analysis of vesicles binding for DNPs of (A) DNP-1, (B) DNP-2, (C) DNP-3, and (D) DNP-4. The data points represent averages and the standard error from at least three independent experiments.

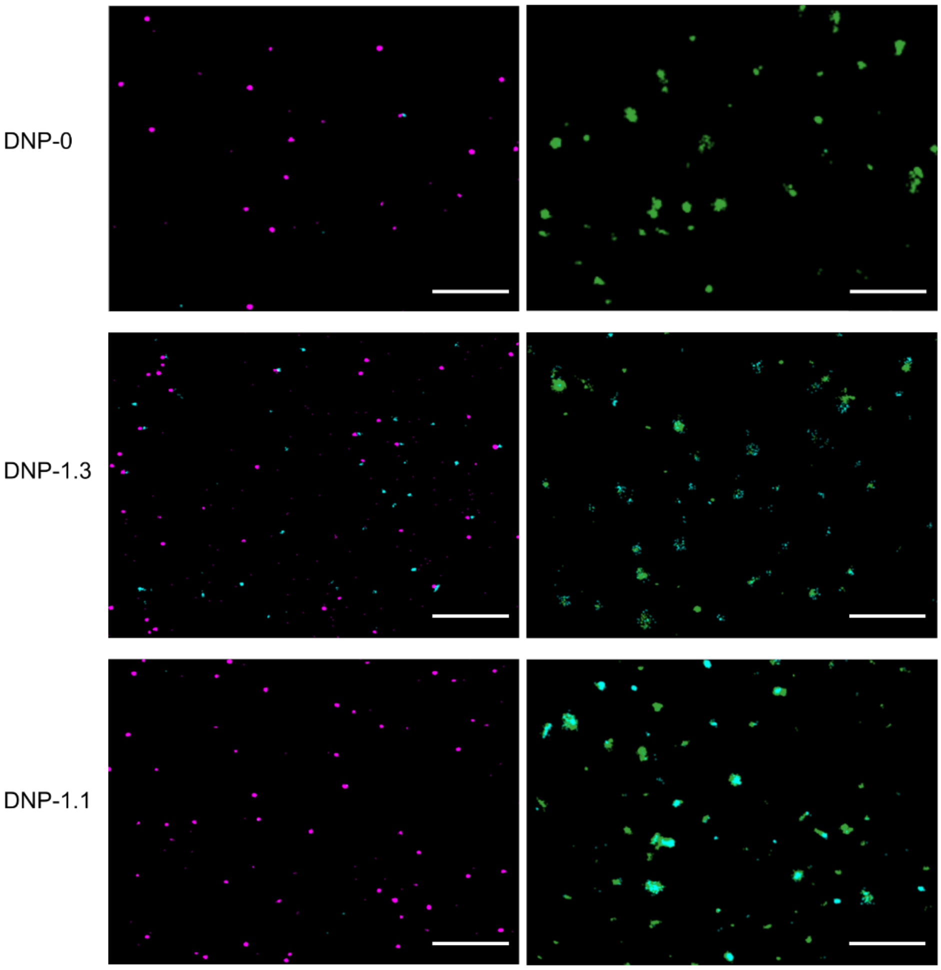


**Figure S16.** dSTORM analysis confirms the size-selective binding of DNA nanoprobe DNP-1.1 to POPC LUVs. dSTORM micrographs of isolated SUVs (left panels) and LUVs (right panels) incubated with DNP-0 without lipid anchor, DNP-1.3 carrying three cholesterol anchors at the corner of the baseplate, and DNP-1.1 with one cholesterol at the corner. DNPs and vesicles were labeled with fluorophores ATTO488 and Cy5, respectively. The fluorophore signals for SUVs are color-coded in magenta, LUVs in green, and DNPs in light blue. The scale bars are 2 μm for both SUVs and LUVs.

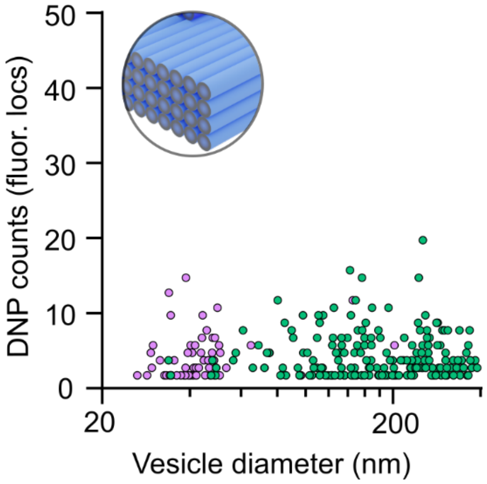


**Figure S17.** Scatter plot on the dSTORM analysis of DNP-0 binding to POPC SUVs and LUVs. Each dot in a scatter plot represents the accumulated DNP fluorescence within a vesicle (vertical axis) as a function of the corresponding vesicle diameter (horizontal axis). Signals from SUVs and LUVs are represented as magenta and green, respectively.


(1) Kim, D. N.; Kilchherr, F.; Dietz, H.; Bathe, M. Quantitative Prediction of 3D Solution Shape and Flexibility of Nucleic Acid Nanostructures. *Nucleic Acids Res.* **2012**, *40*, 2862–2868.

(2) Kučerka, N.; Nieh, M. P.; Katsaras, J. Fluid Phase Lipid Areas and Bilayer Thicknesses of Commonly Used Phosphatidylcholines as a Function of Temperature. *Biochim. Biophys. Acta - Biomembr.* **2011**, *1808*, 2761–2771.
